# Temporally dynamic antagonism between transcription and chromatin compaction controls stochastic photoreceptor specification

**DOI:** 10.1101/2021.08.27.457982

**Authors:** Lukas Voortman, Caitlin Anderson, Elizabeth Urban, Mini Yuan, Sang Tran, Alexandra Neuhaus-Follini, Josh Derrick, Thomas Gregor, Robert J. Johnston

## Abstract

Stochastic mechanisms diversify cell fates during development. How cells randomly choose between two or more fates remains poorly understood. In the *Drosophila* eye, the random mosaic of two R7 photoreceptor subtypes is determined by expression of the transcription factor Spineless (Ss). Here, we investigated how *cis-*regulatory elements and *trans* factors regulate nascent transcriptional activity and chromatin compaction at the *ss* gene locus during R7 development. We find that the *ss* locus is in a compact state in undifferentiated cells. An *early enhancer* drives *ss* transcription in all R7 precursors to open the *ss* locus. In differentiating cells, transcription ceases and the *ss* locus stochastically remains open or compacts. In Ss^ON^ R7s, *ss* is open and competent for activation by a *late enhancer*, whereas in Ss^OFF^ R7s, *ss* is compact and repression prevents expression. Our results suggest that a temporally dynamic antagonism, in which transcription drives decompaction and then compaction represses transcription, controls stochastic cell fate specification.

## Introduction

Cell fate specification is controlled by lineage, signaling, and stochastic regulatory inputs, leading to highly precise developmental outcomes (Petkova et al., 2019). Stochastic mechanisms promote diversity in populations of photoreceptors, olfactory neurons, motor neurons, and immune cells (Alqadah et al., 2016; Bell et al., 2007; Dasen et al., 2003, 2005; Duffy et al., 2012; Johnston and Desplan, 2010; Miyamichi et al., 2005; Ressler et al., 1993; Vassar et al., 1993). Despite the importance of stochastic cell fate specification, how cells randomly choose between two or more fates is poorly understood.

Stochastic cell fate specification is best understood in prokaryotes. One well-characterized example is the bet-hedging mechanism utilized by *Bacillus subtilis*. To minimize losses in a changing environment, populations of genetically identical bacteria maintain a subpopulation of cells that are competent for DNA uptake (Dubnau, 1999; Hahn et al., 1998; Nester and Stocker, 1963). The transient and random transition into the competent fate is controlled by the expression of the transcriptional regulator ComK (Hahn et al., 1998; Turgay et al., 1997). Though most cells maintain low expression of ComK, a subset will experience a pulse of ComK expression that exceeds a threshold and induces a transition to the competent fate (Maamar et al., 2007; Süel et al., 2006). A similar mechanism occurs in the HIV life-cycle, where transcription of the regulatory factor Trans-Activator of Transcription (Tat) determines the switch from proviral latency to active replication (Hendy et al., 2017; Weinberger et al., 2008). Thus, stochastic cell fate specification often requires a pulse of expression of a critical regulator that determines a fate decision.

In addition to transcriptional dynamics, chromatin-mediated repression is a key mechanism mediating stochastic fate specification. In mice, each olfactory sensory neuron (OSN) expresses only one olfactory receptor (OR) gene from a battery of ∼1300 possibilities (Buck and Axel, 1991; Chess et al., 1994; Godfrey et al., 2004). Despite residing in numerous clusters across many chromosomes, all ∼1300 OR genes are repressed and coalesce into heterochromatic foci within the nucleus prior to OR selection (Clowney et al., 2012; Magklara et al., 2011; Sullivan et al., 1996; Zhang and Firestein, 2002). In mutants that impact chromatin modifications and nuclear organization, co-expression of multiple ORs is observed (Clowney et al., 2012). While the mechanism of selection remains elusive, a single OR allele escapes the repressive heterochromatic environment and is expressed in each OSN (Armelin-Correa et al., 2014; Lyons et al., 2013). Thus, chromatin-mediated silencing and selective de-silencing are paramount for the stochastic expression of a single OR gene.

The random mosaic of R7 photoreceptors in the fly eye provides a paradigm to study the integration of transcription and chromatin-mediated repression in stochastic cell fate specification. In the fly eye, stochastic expression of the PAS-bHLH transcription factor Spineless (Ss) in R7 photoreceptors establishes the random pattern of two R7 subtypes across the retina. Ss^ON^ R7s express Rhodopsin 4 (Rh4), while Ss^OFF^ R7s express Rhodopsin 3 (Rh3)(**Fig. 1A-B**)(Bell et al., 2007; Duncan et al., 1998; Johnston and Desplan, 2014; Montell et al., 1987; Wernet et al., 2006) In wild type retinas, 67% of R7s adopt the Ss^ON^/Rh4 fate, while the remaining 33% become Ss^OFF^/Rh3 expressing (**Fig. 1B**). In *ss* protein null mutants, all R7s express Rh3 (**Fig. 1B**). Previously, we showed that stochastic ON/OFF *ss* expression is controlled by an enhancer (*late enhancer, LE)* that is sufficient to drive expression in all R7s and silencers that are required to limit expression to a subset of R7s (**Fig. 1C**)(Johnston and Desplan, 2014).

**Figure 1.**
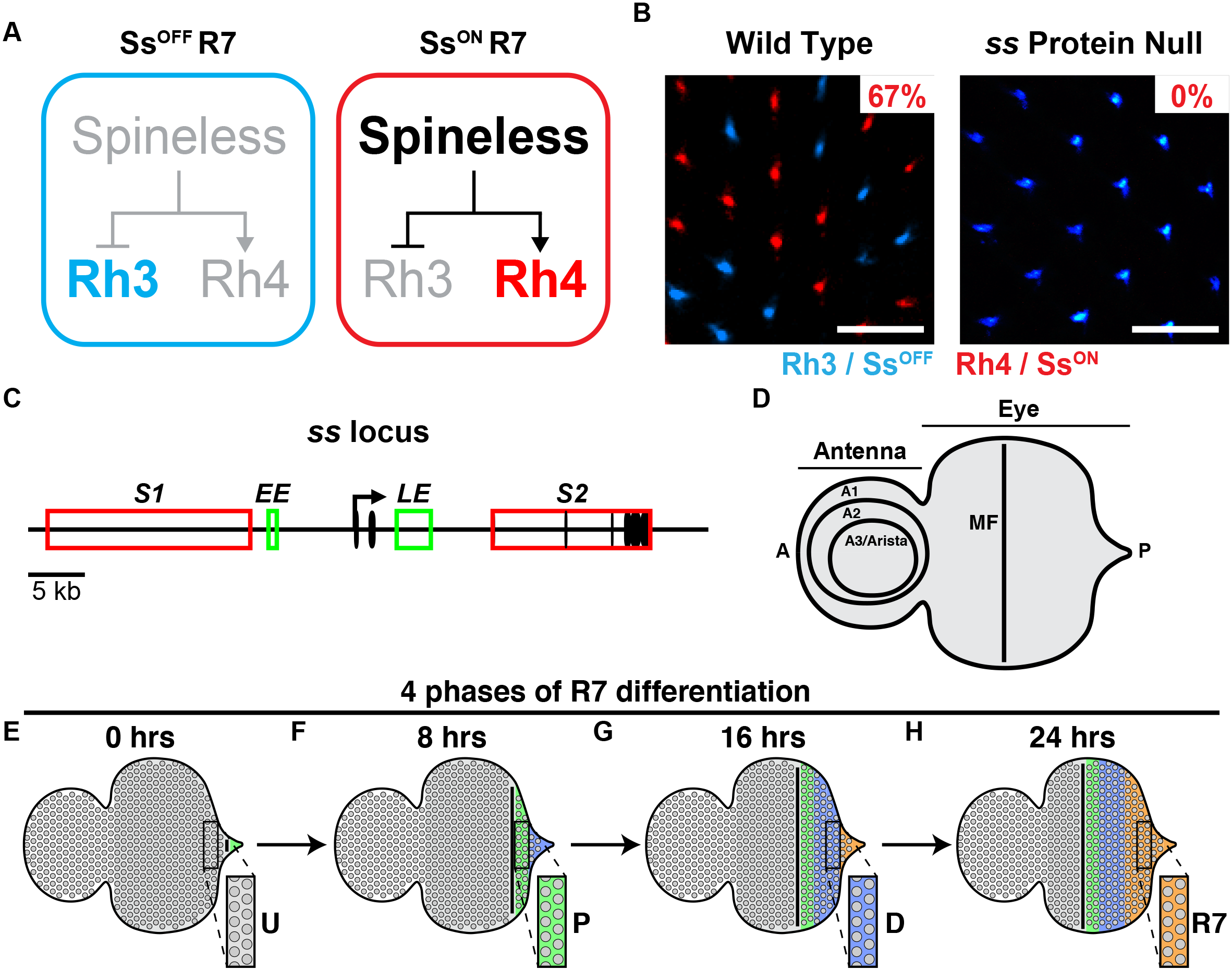
*ss* controls R7 subtype specification. **A)** R7 subtype specification. Expression of Spineless (Ss) promotes the Rh4-expressing R7 fate (red). Absence of Ss yields the Rh3-expressing R7 fate (blue). **B)** Wild type retinas contain 33% Rh3-expressing Ss^OFF^ R7s and 67% Rh4-expressing Ss^ON^ R7s in a random pattern (left). *ss* protein null adult retinas contain only Rh3-expressing Ss^OFF^ R7s and no Rh4-expressing Ss^ON^ R7s (right). Blue = Rh3; red = Rh4. Scalebar = 20 µm. **C)** *ss* gene locus. Black oval = exon; black arrow = promoter; red rectangle = silencer; green rectangle = enhancer; *S1* = *silencer 1*; *S2* = *silencer 2*; *EE* = *early enhancer*; *LE* = *late enhancer*. **D)** Schematized eye-antennal imaginal disc. Antenna is subdivided into the A1, A2, and A3/arista. A = anterior; P = posterior; MF = morphogenetic furrow. **E-H)** Schematized depiction of R7 maturation across time. The magnified inset rectangles illustrate how individual cells proceed through the four phases of development over time. Gray = undifferentiated cells (U); green = precursors (P); blue = differentiating cells (D); orange = R7s (R7).

Here we describe a mechanism that controls stochastic R7 subtype specification in the developing eye. Initially, the *ss* locus is compact in all undifferentiated cells. An *early enhancer* (*EE*) drives *ss* expression that opens the *ss* locus in all R7 precursors during larval development. Expression ceases and the *ss* locus randomly compacts or remains open. In R7s in which *ss* remains open, the *late enhancer* drives *ss* expression and Ss^ON^ R7 fate. In R7s with compact chromatin, repression prevents expression driven by the *late enhancer* yielding the Ss^OFF^ R7 fate. Our data suggest that stochastic fate specification is controlled by the dynamic, intertwined relationship of transcription and chromatin: transcription opens chromatin then chromatin compaction represses transcription. We find that transcription is a source of stochasticity as modulating early transcription in precursors alters the proportions of alternative R7 photoreceptor fates.

## Results

### *ss* expression is dynamic in developing R7s

Photoreceptor identity, including R7 subtype, is specified during larval development in the eye-antennal imaginal disc (**Fig. 1D**). Retinal differentiation begins at the posterior end of the disc and progresses in a wave anteriorly. An indentation in the tissue called the morphogenetic furrow (MF) appears at the posterior end of the disc (**Fig. S1A**). The MF furrow progresses in a developmental wave from posterior to anterior (**Fig. S1**). Behind the MF, photoreceptors differentiate in a stereotypical progression: R8 (**Fig. S1B**), R2/R5 (**Fig. S1C**), R3/R4 (**Fig. S1D**), R1/R6 (**Fig. S1E**), and finally R7 (**Fig. S1F**). As the eye develops in this spatiotemporal manner, analyses of individual discs provides information on all stages of photoreceptor specification, with undifferentiated cells in the anterior and the most differentiated cells in the posterior (**Fig. S1F**)(Ready et al., 1976; Tomlinson and Ready, 1987a, 1987b; Treisman, 2013; Wolff and Ready, 1991).

In these studies, we defined four phases that R7s proceed through during development, including undifferentiated (U), precursor (P), differentiating (D), and differentiated R7 (R7) (**Fig. 1E-H**). In individual late larval discs, we visualized all four phases (**Fig. 1H**). Undifferentiated cells were anterior to the MF (**Fig. 1E-H**). Posterior to the MF, precursors were located at 0-10 µm, differentiating cells were located at 10-30 µm, and R7s were located at >30 µm (**Fig. 1F-H**). In **Fig. 1E-H** and subsequent schematic figures, we diagram only undifferentiated, precursor, differentiating, and R7 cells in the eye and developing cells in the antenna for simplicity.

To characterize *ss* expression in individual cells, we performed nascent RNA fluorescence *in situ* hybridization (RNA FISH). We generated oligo probes covering the entire *ss* transcript, including introns and exons (**Fig. S2A**), and performed single molecule RNA FISH (smFISH)(Beliveau et al., 2012; Little et al., 2013). This strategy yielded single bright fluorescent punctae in *ss-*expressing nuclei, indicating sites of nascent transcription (**Fig. 2A, 2C**). Our observation of one puncta per nucleus is consistent with chromosome pairing in close proximity in somatic cells of *Drosophila* (Stevens, 1908). This approach enabled quantification of *ss* transcription in each developmental context.

**Figure 2.**
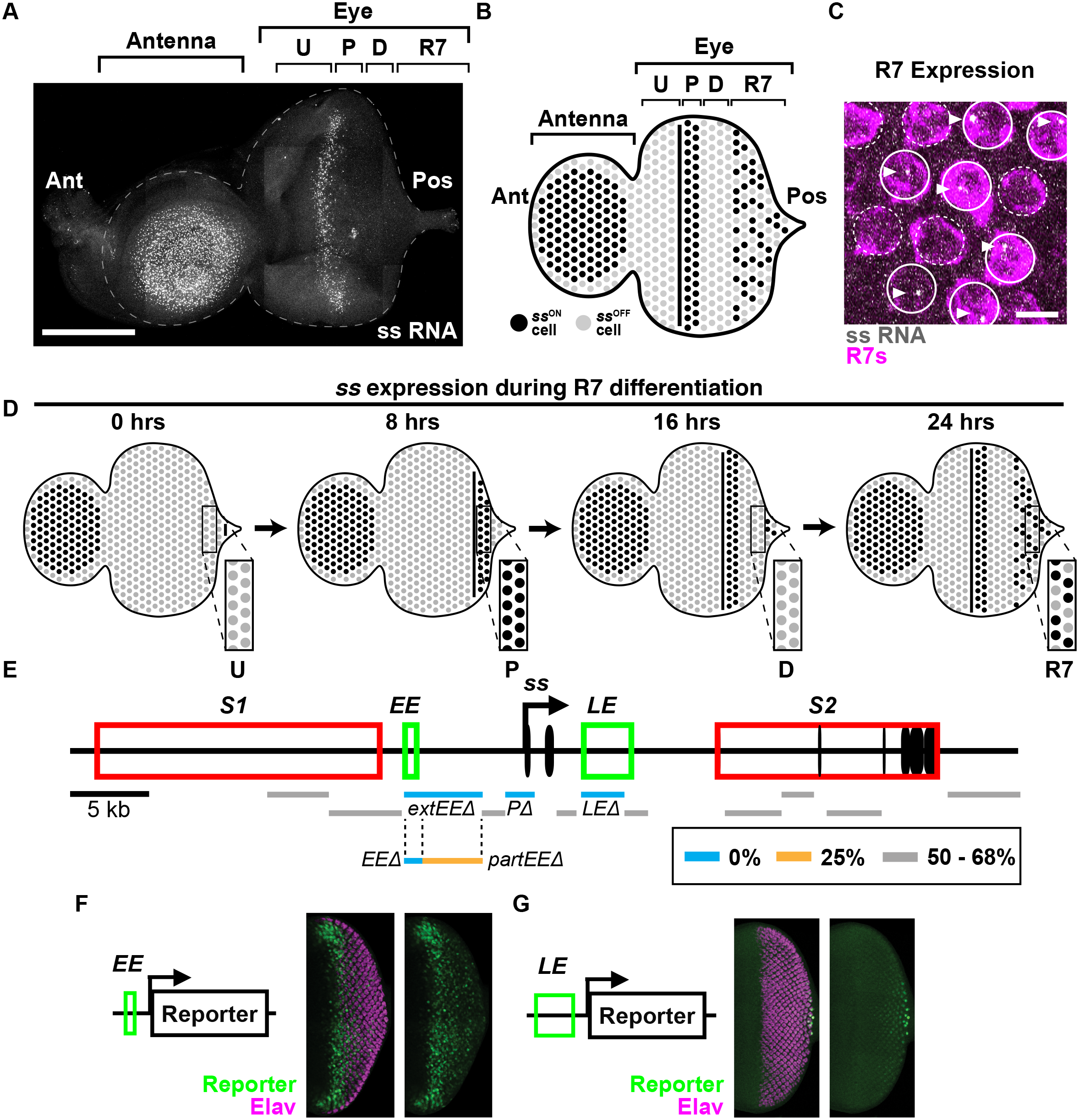
Two temporally distinct enhancers drive *ss* expression in the developing eye. **For A-B, D)** U = undifferentiated cells; P = precursors; D = differentiating cells; R7 = R7s. Ant = anterior; Pos = posterior. **A)** *ss* RNA is expressed in antenna cells, precursors, and R7s in the larval eye-antennal imaginal disc. Gray = *ss* RNA. Scalebar = 100 µm. **B)** Schematized eye antennal-disc imaginal disc. *ss* expression as in A. **C)** Nascent *ss* RNA transcripts in a subset of R7s in the larval eye. Magenta = R7 reporter; gray = *ss* RNA; sold line circles = Ss^ON^ R7s, dashed line circles = Ss^OFF^ R7s. Scalebar = 5 µm. **D)** Schematized depiction of *ss* expression across time. The magnified inset rectangles illustrate *ss* expression dynamics in individual cells during the four phases of development over time. **E)** *ss* gene locus and CRISPR deletion screen. Deletion of *extEE, EE, LE,* and *P* displayed 0% Ss^ON^ R7s. Deletion of *partEE* displayed 25% Ss^ON^ R7s. Black oval = exon; black arrow = promoter; red rectangle = silencer; green rectangle = enhancer; blue line = CRISPR deletion with major phenotypic effect (0% Ss^ON^ R7s); orange line = CRISPR deletion with partial phenotypic effect (25% Ss^ON^ R7s); gray line = CRISPR deletion with minimal phenotypic effect (50-68% Ss^ON^ R7s); *S1* = silencer 1; *S2* = silencer 2; *EE* = early enhancer; *LE* = late enhancer. **For F, G)** Green = reporter; magenta = Elav (neurons). **F)** The *early enhancer* reporter is expressed in precursors. **G)** The *late enhancer* reporter is expressed in R7s.

*ss* is strongly expressed in the antennal region of the eye-antennal disc, serving as a positive control (**Fig. 2A**). *ss* is not expressed in the peripodial membrane that overlies the eye-antennal disc, acting as a negative control (**Fig. S2B**).

In the eye, *ss* is differentially expressed during the progressive temporal phases of R7 specification. *ss* is not expressed in undifferentiated cells, *ss* is strongly expressed in all precursors, *ss* is not expressed in differentiating cells, and *ss* is expressed in a subset of R7s (**Fig. 2A-D, Fig. S2C**). The *ss*^ON/OFF^ ratio in larval R7s is similar to the *ss*^ON/OFF^ ratio in adult R7s (**Fig. S2D**), consistent with this decision being made in larvae and maintained throughout the lifetime of the organism (Johnston and Desplan, 2014). As expression in an individual disc represents different temporal phases of R7 development, we conclude that *ss* expression is dynamic as R7s develop: off in the undifferentiated cell phase, on in the precursor phase, off in the differentiating cell phase, and finally, on in a subset of R7s in the differentiated R7 phase (**Fig. 2A-D**).

### Two temporally distinct enhancers drive *ss* expression in the developing eye

To identify *cis*-regulatory elements that regulate stochastic ON/OFF expression of *ss* in R7s, we used CRISPR to make a series of 1-5 kb deletions in the *ss* locus (**Fig. 2E, Table S1**). As Rhodopsin expression faithfully reports Ss expression state in adults (Rh4 = Ss^ON^, Rh3 = Ss^OFF^)(Johnston and Desplan, 2014), we examined Rh3 and Rh4 expression and determined the proportions of R7 subtypes. We identified three elements that are required for *ss* expression in R7s, including the promoter (*P*), a 5.4 kb upstream element (*extEE*), and a previously identified 3 kb intronic R7/R8 enhancer (*LE*) (**Fig. 2E, Table S1**)(Johnston and Desplan, 2014). Deletion of these regions reduced the proportion of Ss^ON^/Rh4 R7s to 0% (**Fig. 2E, Table S1**). Therefore, these cis-regulatory regions are required for normal *ss* expression.

We conducted additional partial deletions of the *extEE* region to determine a minimal *cis*-regulatory region required for *ss* expression. Deletion of the 1.3 kb *EE* region caused a dramatic decrease of Ss^ON^/Rh4 R7s to 0% (**Fig. 2E, Table S1**), while deletion of the neighboring 4.1 kb *partEE* caused a partial reduction of Ss^ON^/Rh4 R7s to 25% (**Fig. 2E, Table S1**). As *EE* was strictly required for *ss* expression, we interrogated this region further.

To assess the spatiotemporality of *EE* and *LE* activities, we generated enhancer reporter constructs and examined expression in larval eye-antennal discs. The *EE* element drove expression in precursors similar to *ss* RNA expression (**Fig. 2F**). In contrast, the *LE* intronic element drove expression in all R7s (**Fig. 2G**)(Johnston and Desplan, 2014). Thus, *EE* and *LE* are sufficient to drive expression in precursors and R7s respectively.

As chromatin accessibility is associated with enhancer activity, ATACseq experiments can predict candidate enhancers (Buenrostro et al., 2015). We analyzed published scATACseq datasets from developing fly eye-antennal discs (**Fig. S2E**)(Bravo González-Blas et al., 2020). For antennal cells that express *ss*, chromatin accessibility peaks were observed at the promoter, but not at *EE* or *LE* (**Fig. S2E**). For precursors that express *ss*, peaks occurred at the *EE* and *promoter*, but not *LE* (**Fig. S2E**). For all PRs, of which only a subset of R7s express *ss,* peaks were observed at the *LE* and *promoter* and were significantly reduced for the *EE* (**Fig. S2E**). As a small peak remains at the *EE* in R7s, some residual chromatin accessibility may remain at this later timepoint. Alternatively, some cells may have been incorrectly clustered into this cell type. These observations support roles for the *EE* and *LE* as enhancers that drive *ss* expression during distinct temporal phases of R7 development.

Together, these data suggest that *ss* expression in the eye is controlled by two temporally distinct enhancers, one that drives early expression in precursors (*early enhancer*, *EE*) and one that drives late expression in R7s (*late enhancer*, *LE*). We next sought to determine the role for temporally distinct *ss* expression in R7 development.

### Early *ss* expression in precursors is required for *ss* expression in R7s

To test how the *early* and *late enhancers* regulate *ss* expression during photoreceptor development, we observed *ss* expression in mutant conditions. In the developing fly eye, automated identification and assignment of nascent spots to individual cells is challenging in three-dimensions and not necessary to describe the changes in expression observed here. Therefore, to quantify *ss* expression in the eye-antennal disc, we measured the density of nascent RNA spots per unit area (µm^2^) (**Fig. S3A**) (see **Methods**). We assessed Rh4/Ss^ON^ and Rh3/Ss^OFF^ in R7s in adult retinas (**Fig. 3D, W**). *ss* expression in antennal cells in larval eye-antennal discs served as a positive internal control (**Fig. 3B, U**). Promoter deletion (*PΔ*) mutants acted as a negative control, exhibiting a complete loss of *ss* expression in antennal cells, precursors, and R7s (**Fig. 3E-H, U-W**).

**Figure 3.**
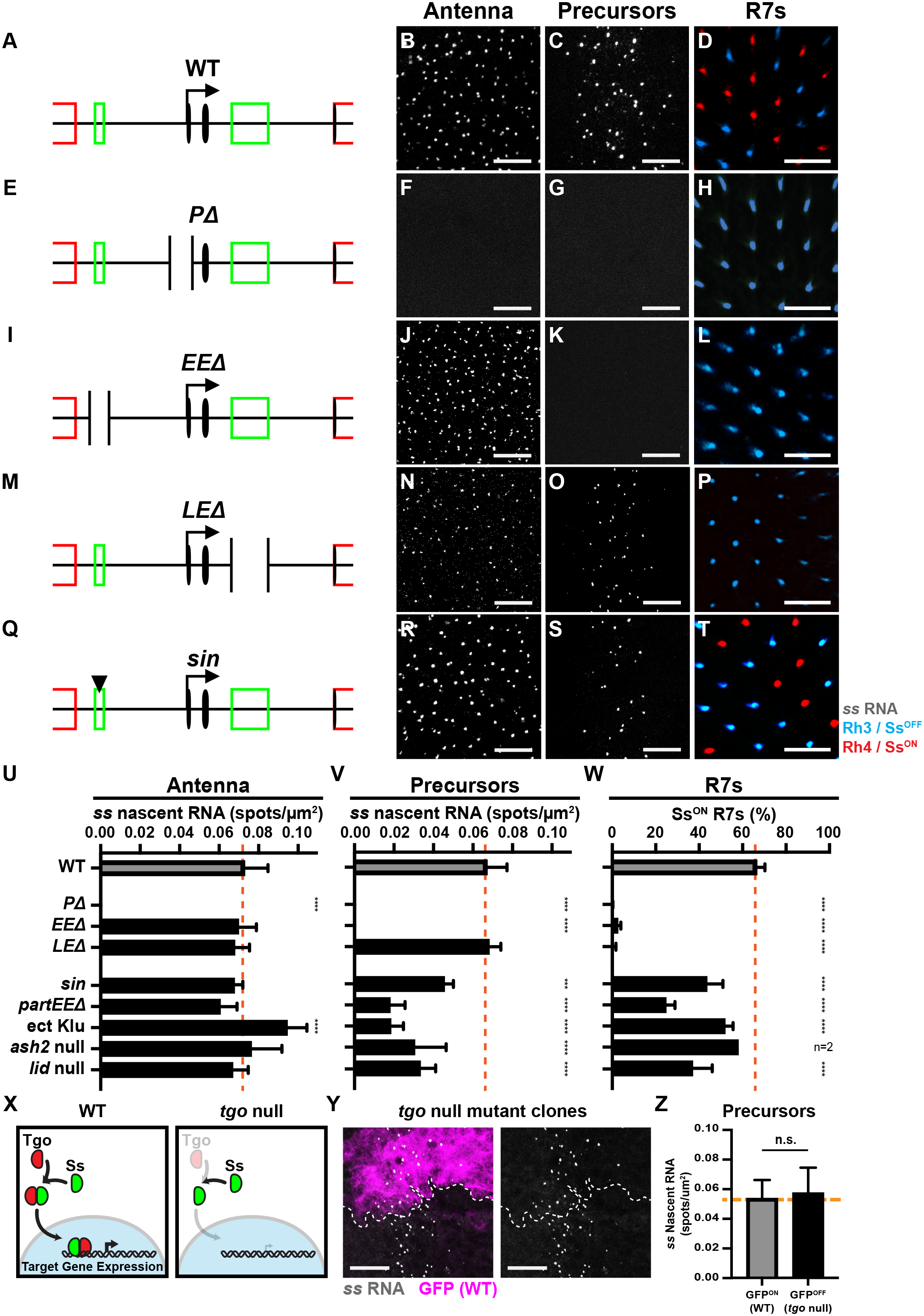
Decreasing early *ss* expression decreases the proportion of Ss^ON^ R7s. For A, E, I, M, Q) Truncated schematized *ss* locus. **For B, F, J, N, R)** *ss* RNA expression in the antenna. Gray = *ss* RNA. Scalebar = 10 µm. **For C, G, K, O, S)** Early *ss* RNA expression in precursors. Gray = *ss* RNA. Scalebar = 10 µm. **For D, H, L, P, T)** Adult Rh3/Ss^OFF^ and Rh4/Ss^ON^ expression in R7s. Blue = Rh3; red = Rh4. Scalebar = 20 µm. **A-D)** In wild type (WT) animals, *ss* is expressed in antennal cells, precursors, and a subset of R7s. **E-H)** In *promoter* (*PD*) mutants, *ss* is not expressed in antennal cells, precursors, or R7s. **I-L)** In *early enhancer* (*EED*) mutants, *ss* is expressed in antennal cells but not expressed in precursors or R7s. **M-P)** In *late enhancer* (*LED*) mutants, *ss* is expressed in antennal cells and precursors, but not expressed in R7s. **Q-T)** Animals with a high affinity Klu repressor binding site (*sin*) in the *early enhancer* display normal *ss* expression in antennal cells, reduced *ss* expression in precursors, and a decrease in % Rh4/Ss^ON^ R7s. **For U-W, Z)** Orange line indicates mean WT expression. n.s. denotes p > 0.05; *** denotes p < 0.0005; **** denotes p < 0.0001. **U)** Quantification of *ss* expression in antennal cells for **B, F, J, N, R**. **V)** Quantification of *ss* expression in precursors for **C, G, K, O, S**. **W)** Quantification of % Rh4/Ss^ON^ R7s for **D, H, L, P, T**. **X)** Cartoon depicting the Ss/Tgo mechanism of action in WT and its breakdown in *tgo* null mutants. Red = Tgo; green = Ss; blue = nucleus. **Y)** *ss* RNA is expressed in precursors in *tgo* null mutant clones. Gray = *ss* RNA; magenta = GFP; dashed line = clone boundary. GFP-indicates *tgo* null mutant clone; GFP+ indicates wild type clone. Gray = *ss* RNA; magenta = GFP; dashed line = clone boundary. Scalebar = 10 µm. **Z)** Quantification of *ss* expression in precursors for **Y**.

We found that *EE*Δ mutants lost *ss* expression in precursors and Rh4/Ss^ON^ in R7s, while *ss* expression was maintained in antennal cells (**Fig. 3I-L, U-W**). *LE*Δ mutants displayed a complete loss of Rh4/Ss^ON^ expression in R7s but showed normal *ss* expression in precursors and antennal cells (**Fig. 3M-P, U-W**). Together, these results show that the early expression of *ss* in precursors driven by the *early enhancer* is required for activation by the *late enhancer* in R7s.

### *ss* expression does not require Ss protein feedback in precursors

Early expression often affects later expression from the same gene locus through protein feedback. For example, protein feedback reinforces transcriptional pulses during the stochastic expression of ComK in *Bacillus subtilus* (Maamar et al., 2007; Süel et al., 2006). In the developing fly eye, no detectable Ss protein is made during the early expression in precursors (Johnston and Desplan, 2014). Nevertheless, extremely low levels of Ss protein could trigger regulatory feedback. Ss protein requires heterodimerization with another PAS-bHLH transcription factor, Tango (Tgo) to enter the nucleus and regulate gene expression (**Fig. 3X**)(Thanawala et al., 2013). To test whether Ss/Tgo feedback activity affects early *ss* expression, we generated *tgo* null mutant clones and observed no effect on *ss* transcription in precursors (**Fig. 3Y-Z**). This result suggests that *ss* regulation in the eye does not require Ss protein feedback, consistent with our previous findings (Johnston and Desplan, 2014). This finding also suggests that the early transcription of *ss* activates late expression by a Ss/Tgo-protein independent mechanism.

### Modulating early *ss* expression alters the proportion of Ss^ON^ R7s

The *early enhancer* is required for specification of Ss^ON^ R7s, as knocking out the *early enhancer* caused a complete loss of *ss* expression in R7s (**Fig. 3I-L, U-W**). We hypothesized that reducing activation by the *early enhancer* would decrease the number of *ss* expressing precursors and the proportion of Ss^ON^ R7s.

The *early enhancer* contains a binding site for the transcriptional repressor, Klumpfuss (Klu)(**Fig. S3B**)(Anderson et al., 2017). A single base insertion (“*sin*”) within the *early enhancer* increases the binding affinity of Klu (Anderson et al., 2017). Flies with *sin* displayed a reduction in the number of *ss-*expressing precursors and a decrease in the proportion of Ss^ON^ R7s (45% Ss^ON^) (**Fig. 3Q, S-T, V-W**). Flies with *sin* had no change in *ss* expression in the antenna (**Fig. 3R, U**). Ectopic expression of Klu reduced *ss* expression in precursors and the ratio of Ss^ON^ R7s (51.8% Ss^ON^/Rh4)(**Fig. 3V-W, S3L-M**) (Anderson et al., 2017). Ectopic expression of Klu in precursors caused an increase of *ss* expression in the antenna (**Fig. 3U, S3K**), consistent with differential regulation of *ss* by Klu across tissues through different enhancers (Klein and Campos-Ortega, 1997; Yang et al., 1997). Additionally, a partial deletion of the *early enhancer* (*pEE*Δ), which resulted in a reduction of Ss^ON^ R7s to 25%, also had a reduction in expression in precursors (**Fig. 3U-W, S3H-J**). This deletion removes the sequence abutting the *early enhancer* and may disrupt the binding of other *trans-*acting factors to the *early enhancer*. These data suggest that decreasing *early enhancer* activity by genetically altering *cis* or *trans* inputs reduces *ss* expression in precursors and leads to a reduction in Ss^ON^ R7s.

To identify regulators of R7 subtype specification, we screened flies with mutations or RNAi knockdowns in genes encoding chromatin modifiers for changes in the ratio of Ss^ON^ and Ss^OFF^ R7s (**Table S2**). Reducing activity of two genes encoding chromatin modifiers, *ash2* and *lid*, caused significant loss of Ss^ON^ R7s. Knockdown of the trithorax group gene *ash2* (Adamson and Shearn, 1996; Papoulas et al., 1998) caused a decrease in Ss^ON^ R7s in two independent RNAi lines (**Table S2**). *ash2^1^* null mutants displayed a reduction in Ss^ON^ R7s (38.4%) (**Fig. 3W, S3P**). *ash2^1^* null mutants displayed a cell autonomous decrease in *ss* expression in precursors (**Fig. 3V, S3C, O**), and no change in *ss* expression in antennal cells (**Fig. 3U, S3N**). Similarly, a null mutation in the histone demethylase gene *lid* (Eissenberg et al., 2007; Secombe et al., 2007) caused a reduction in *ss* expression in precursors and the proportion of Ss^ON^ R7s (36.8% Ss^ON^/Rh4), but had no effect on *ss* expression in antennal cells (**Fig. 3U-W, S3Q-S**). These data implicate a role for chromatin modifiers in *ss* regulation and support the conclusion that decreasing *ss* expression in precursors ultimately decreases the proportion of Ss^ON^ R7s.

### Derepression of early *ss* expression increases the proportion of Ss^ON^ R7s

We next investigated how derepression of *ss* affected R7 subtype specification. We hypothesized that mutant genotypes resulting in an increase in Ss^ON^ R7s will also have an altered expression pattern earlier in development. For these experiments, we examined *ss* expression in undifferentiated cells (*ss*^OFF^), precursors (*ss*^ON^), and differentiating cells (*ss*^OFF^) in larval eye-antennal discs as well as R7s (mix of *ss*^ON^ and *ss*^OFF^) in adult retinas (**Fig. 4C-F**).

**Figure 4.**
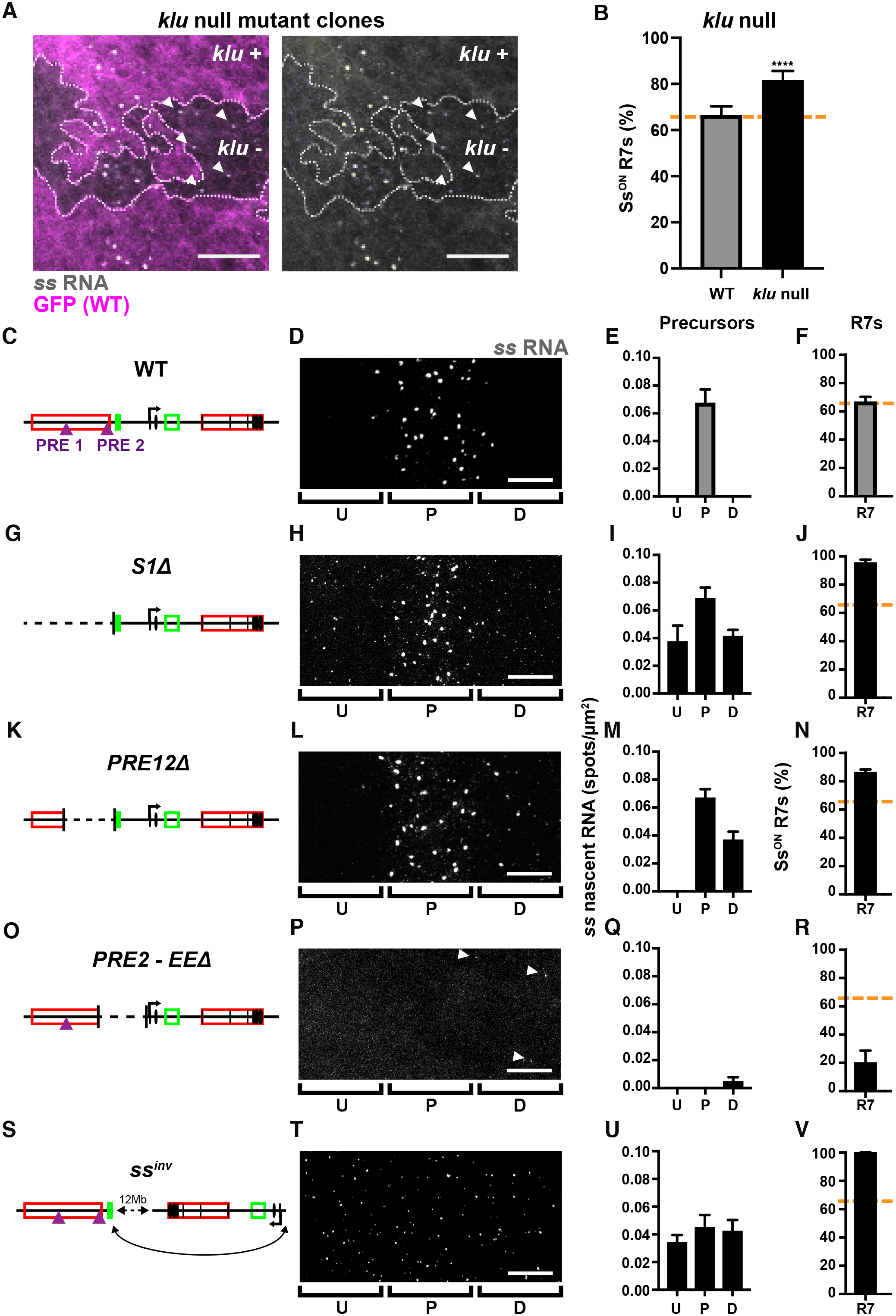
Derepressing early *ss* expression increases the proportion of Ss^ON^ R7s. **A)** Early *ss* expression in precursors is extended into differentiating cells in *klu* null mutant clones. GFP-indicates *klu* null mutant clone; GFP+ indicates wild type clone. Gray = *ss* RNA; magenta = GFP; dashed line = clone boundary; arrows = nascent *ss* RNA in differentiating cells. Scalebar = 10 µm. **For B, F, J, N, R, V)** Orange line indicates mean WT *ss* expression. **For C-V)** U = undifferentiated cells; P = precursors; D = differentiating cells; R7 = R7s. **B)** % Rh4/Ss^ON^ R7s increases in *klu* null mutants. **** denotes p < 0.0001. **For C, G, K, O, S)** Schematized *ss* locus. **For D, H, L, P, T)** *ss* expression in undifferentiated cells, precursors, and differentiating cells. Gray = *ss* RNA. Scalebar = 10 µm. **For E, I, M, Q, U)** Quantification of expression for **D, H, L, P, T**. **For F, J, N, R, V)** Quantification of % Rh4/Ss^ON^ R7s. **C-F)** Wild type (WT) animals display no *ss* expression in undifferentiated cells, expression in precursors, no expression in differentiating cells, and 67% Rh4/Ss^ON^ R7s. **G-J)** *Silencer 1* (*S1D*) mutants display *ss* expression in undifferentiated cells, precursors, and differentiating cells, and 95% Rh4/Ss^ON^ R7s. **K-N)** *PRE1* and *PRE2* (*PRE12D*) double mutants display no *ss* expression in undifferentiated cells, expression in precursors and differentiating cells, and 86% Rh4/Ss^ON^ R7s. **O-R)** *PRE2* and *EE* (*PRE2-EED*) double mutants display no *ss* expression in undifferentiated cells and precursors, expression in differentiating cells, and 20% Rh4/Ss^ON^ R7s. **S-V)** *ss inversion* (*ss^inv^*) mutants display low *ss* expression in undifferentiated cells, precursors, and differentiating cells, and 100% Rh4/Ss^ON^ R7s. See **Figure S4A-E** for statistical comparison between genotypes.

Increasing the binding affinity of a Klu site or increasing Klu levels reduced early *ss* expression and the proportion of Ss^ON^ R7s (**Fig. 3U-W**). In contrast, *klu* null mutant clones displayed a temporal extension of *ss* expression beyond the precursor state into the differentiating state, when *ss* is not normally expressed (**Fig. 4A**). *klu* null mutants also exhibited an increase in the proportion of Ss^ON^ R7s (82% Ss^ON^)(**Fig. 4B**) (Anderson et al., 2017). These data suggest that Klu is a cell-autonomous off switch for *ss* expression and that extended expression of *ss* leads to an increase in the probability of Ss^ON^ R7 fate.

Repressive *silencer* elements restrict expression of *ss* to a subset of R7s (Johnston and Desplan, 2014). We focused on the effects of a 36.4 kb deletion of *silencer1* (*S1*Δ)(**Fig. 4G**)(Thanawala et al., 2013). Heterozygous *S1*Δ/+ mutants displayed *ss* expression in undifferentiated cells prior to the precursor stage and in differentiating cells after the precursor stage. They also displayed an increase in the proportion of Ss^ON^ R7s (95%)(**Fig. 4H-J**)(Johnston and Desplan, 2014). Additionally, *S1*Δ/+ mutants showed low level *ss* expression in most cells of the eye-antennal disc, including the peripodial membrane (*ss*^OFF^ control). Together, these data indicate that *silencer1* is generally required for repression of *ss* (**Fig. S4F**).

As *ss* expression was diminished in *ash2* and *lid* mutants, we hypothesized that chromatin is playing a role in *ss* repression. We examined the region deleted in *S1*Δ mutants for Polycomb response elements (PREs). PREs are DNA elements bound by Polycomb group (PcG) proteins that nucleate repressive heterochromatin (Chan et al., 1994; Paro and Hogness, 1991; Simon et al., 1993; Strutt et al., 1997). ChIP-seq experiments in whole embryos and larvae showed distinct ChIP peaks for Polycomb group (PcG) proteins including Polycomb (Pc), Posterior sex combs (Psc), and Sex combs extra (dRing), suggesting several putative PREs in the *ss* locus (Celniker et al., 2009). Two putative PREs fall within the region deleted in the *S1*Δ mutants (**Fig. S4H**). We generated a deletion that removed 13 kb containing both PREs (*PRE12*Δ)(**Fig. 4K**). Hemizygous *PRE12*Δ mutants displayed a temporal extension of *ss* expression into differentiating cells and an increase in the ratio of Ss^ON^ R7s (86%) but did not exhibit ectopic expression in undifferentiated cells (**Fig. 4L-N**). These data suggest that the two PREs repress *ss* expression. Together, chromatin regulation at the *ss* locus is critical for R7 subtype specification and extending early expression increases the proportion of Ss^ON^ R7s.

### Derepression restores Ss^ON^ R7 fate in *early enhancer* mutants

Our data suggest that activation in precursors precedes repression in differentiating cells during stochastic R7 subtype specification. To test the temporality of these steps, we examined mutants that impaired activation in precursors and repression in differentiating cells. We predicted that derepression in differentiating cells would offset loss of activation in precursors to restore Ss^ON^ R7 fate. We used imprecise P-element excision to generate an 11.8 kb mutant that deleted one PRE within *silencer1* and the *early enhancer* (*PRE2-EE*Δ)(**Fig. 4O**). In *PRE2-EE*Δ mutants, *ss* expression in precursors was completely lost (**Fig. 4P-Q**), consistent with the loss of activation by the *early enhancer*. *PRE2-EE*Δ mutants displayed low levels of *ss* expression in differentiating cells (**Fig. 4P-Q**), consistent with a loss of repression. *PRE2-EE*Δ mutants contained 20% Ss^ON^ R7s (**Fig. 4R**), suggesting that depression in differentiating cells restores Ss^ON^ R7 fate in the absence of activation by the *early enhancer*.

To further test these interactions, we examined flies with an inversion (*ss^inv^*) that moves the *ss* promoter and *late enhancer* ∼12 Mb away from upstream regions, preventing regulation by the *early enhancer* and *silencer1* (**Fig. 4S-V, S4G**)(Thanawala et al., 2013). *ss^inv^* mutants displayed weak *ss* expression in undifferentiated cells, precursors, and differentiated cells (**Fig. 4T-U**), consistent with the loss of repression by *silencer1*. Strong *ss* expression in precursors was decreased (**Fig. 4T-U**), consistent with the loss of activation by the *early enhancer*. *ss^inv^* mutants contained 100% Ss^ON^ R7s (**Fig. 4V**), suggesting that depression enables Ss^ON^ R7 fate when activation by the *early enhancer* is lost. These data are consistent with activation driven by the *EE* in precursors before repression by *silencer* elements in differentiating cells during stochastic R7 subtype specification.

### Repression limits *ss* expression to a subset of R7s

To directly test the hypothesis that repression limits *ss* expression to Ss^ON^ R7s, we developed a “repression reporter” strategy. When inserted into a control locus on the X chromosome, a broad photoreceptor enhancer reporter (*[3xP3]>RFP*)(**Fig. 5A**)(Bischof et al., 2007) drives expression in all photoreceptors in all ommatidia of the adult retina (**Fig. 5B-C, F**). We used CRISPR to insert this reporter into different locations in the endogenous *ss* locus (**Fig. 5A,** inserts 1-4). We hypothesized that, if the local chromatin environment of the *ss* locus was sufficient to repress expression, expression of the reporter would be limited to Ss^ON^ R7s. If the *ss* locus did not repress expression, the reporter would be expressed in all photoreceptors, including all R7s.

**Figure 5.**
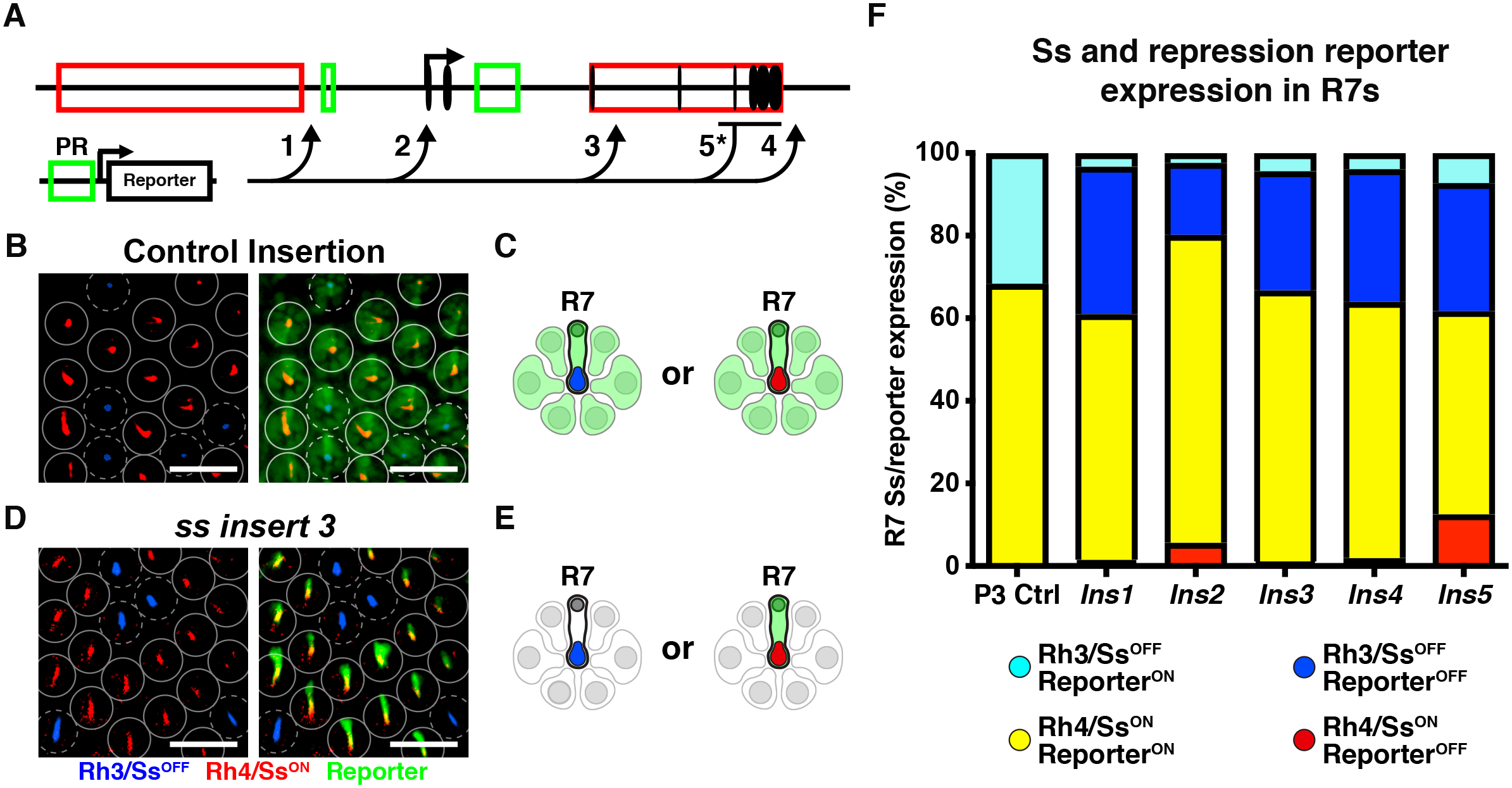
Repression by the *ss* locus limits expression to a subset of R7s. **A)** Schematic of the broad photoreceptor (PR) enhancer reporter construct and insertion sites into the *ss* locus. Arrows = insertion sites. 1-4 were inserted using CRISPR. 5* was inserted using homologous recombination to replace part of the locus. **For B-E)** Blue = Rh3/Ss^OFF^ R7s; red = Rh4/Ss^ON^ R7s; green = reporter. **B)** When inserted at a control site, the PR enhancer reporter is expressed in all PRs, including Rh3/Ss^OFF^ R7s and Rh4/Ss^ON^ R7s. Scalebar = 20 µm. **C)** Cartoon depiction of **B**. **D)** When inserted into the *ss* locus, the PR enhancer reporter is expressed in Rh4/Ss^ON^ R7s but is not expressed in Rh3/Ss^OFF^ R7s or outer PRs. Scalebar = 20 µm. **E)** Cartoon depiction of **D**. **F)** Quantification of rhodopsin and reporter expression. Cyan = Rh3/Ss^OFF^, reporter^ON^; blue = Rh3/Ss^OFF^, reporter^OFF^; yellow = Rh4/Ss^ON^, reporter^ON^; red = Rh4/Ss^ON^, reporter^OFF^.

When inserted into four different locations in the *ss* locus, the reporter transgene was nearly perfectly repressed in Ss^OFF^ R7s and expressed in Ss^ON^ R7s in adult retinas (**Fig. 5D-F**). Ss^ON^ R7s had greater than 93% co-expression with the reporter, while Ss^OFF^ R7s expressed the reporter in less than 14% of cells (**Fig. S5A**). With the exception of *ins2*, inserted into the 5’ UTR of *ss*, the reporter lines did not significantly affect the ratio of Ss^ON/OFF^ R7s (**Fig. S5B**). Expression of the reporter in Ss^ON^ R7s and repression in Ss^OFF^ R7s suggest that repression at the *ss* locus limits expression to a subset of R7s.

When inserted into control loci, the reporter also drives expression in the motion-detecting outer photoreceptors (R1-6) (**Fig. 5B-C**). When inserted into the *ss* locus, the reporter was not expressed in outer photoreceptors (**Fig. 5D-E**), suggesting that the *ss* locus represses expression in these cells.

In control insertion lines, the reporter also drives expression in color-detecting R8 photoreceptors. When inserted into the *ss* locus, the reporter remained expressed in a subset of R8s (**Fig. S5C**). In these lines, Ss was expressed in the same subset of R8s (**Fig. S5C**). These observations suggest that the broad photoreceptor enhancer reporter, inserted into the *ss* locus, was sufficient to ectopically drive *ss* and reporter expression, overcoming repression in R8s.

Regulation of the “repression reporter” may be specific to this enhancer reporter construct inserted by CRISPR. We previously used homologous recombination to replace the last four exons of *ss* with a different broad photoreceptor enhancer reporter (*GMR>GFP*, **Fig. 5A, insert 5**)(Thanawala et al., 2013). For this reporter, we examined *ss^Ins5^/+* heterozygous flies. Similar to the other reporter, expression of this reporter was generally expressed in Ss^ON^ R7s (79.5% co-expressing), repressed in Ss^OFF^ R7s (18.7% expressing), repressed in outer photoreceptors, and expressed in R8s (**Fig. 5F, S5A**).

Insertion of two different types of broad photoreceptor enhancer reporters by two different methods across five locations in the *ss* gene locus resulted in repression of the reporter in Ss^OFF^ R7s and expression in Ss^ON^ R7s. These data indicate that the *ss* gene locus represses expression and suggests a role for the local chromatin environment in repression.

### Visualizing chromatin compaction at the *ss* locus

The repression reporter strategy showed that the *ss* locus restricts expression to a subset of R7s, likely through chromatin remodeling of the *ss* locus during R7 subtype specification. Additionally, two silencer elements are required for proper *ss* expression, consistent with a role for long range repressive interactions possibly through chromatin compaction.

We sought to characterize the compaction state of the *ss* gene locus during R7 subtype specification. The heterogeneity of cell types in the larval fly eye and limiting quantities of cells impede cell-type-specific analyses through ChIP-seq and ATAC-seq approaches. To examine chromatin compaction with single cell resolution in intact tissue, we developed a 3-color DNA-FISH strategy. We labeled a 50 kb upstream region, a 65 kb region encompassing *ss,* and a 50 kb downstream region with different fluorescently labeled probes (**Fig. 6A**). We identified the 3-D center of the spheroid for each region and measured the total 3-D distance from the upstream region to the *ss* region, and from the *ss* region to the downstream region in individual nuclei (**Fig. 6B-C**)(Joyce et al., 2012; Rosin et al., 2018; Viets et al., 2019). Larger distances reflect a more open state, while smaller distances indicate a more compact state (**Fig. 6B-C**). We hypothesized that transcribed *ss* loci would be more open compared to inactive *ss* loci which would be more compact.

**Figure 6.**
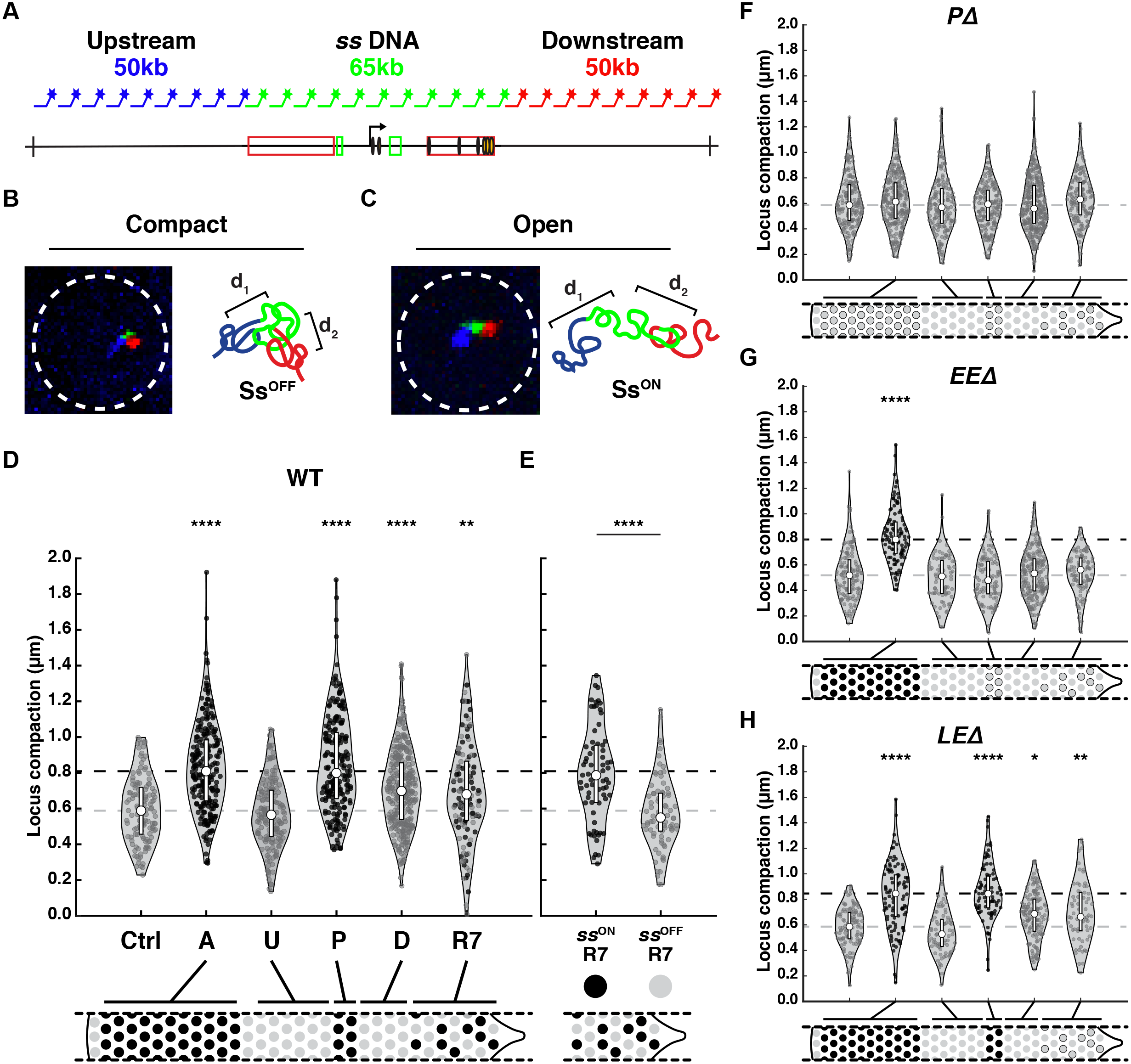
Dynamic chromatin compaction of the *ss* locus is dependent on early transcription in precursors. **For A-C)** Blue = *ss* upstream DNA; green = *ss* locus DNA; red = *ss* downstream DNA. **A)** Schematic of DNA FISH probes used to label the upstream, *ss* locus, and downstream regions of DNA. **B)** Ss^OFF^ cell with compact chromatin at the *ss* locus. Left = image; right = schematized model. **C)** Ss^ON^ cell with open chromatin at the *ss* locus. Left = image; right = schematized model. **For D-H)** Quantification of compaction. Ctrl = peripodial membrane cells; A = antennal cells; U = undifferentiated cells; P = precursors; D = differentiating cells; R7 = R7s. Black circle = *ss*^ON^ cell; gray circle = *ss*^OFF^ cell; white rectangle = quartile; white circle = median; gray dashed line = *ss*^OFF^ control/peripodial membrane median; black dashed line = *ss*^ON^ control/antennal cells median. * denotes p < 0.05; ** denotes p < 0.005; **** denotes p < 0.0001. **For D, F-H,** Ctrl cells were compared to A, U, P, D, or R7 cells. For **F,** Ss^ON^ R7s were compared to Ss^OFF^ R7s. **D)** WT flies. **E)** For WT flies, compaction in Ss^ON^ R7s and Ss^OFF^ R7s labeled by the *Ins5* reporter. **F)** *PD* mutants. **G)** *EED* mutants. **H)** *LED* mutants.

To test our method, we examined compaction in peripodial membrane cells and antennal cells. The *ss* locus was more compact in peripodial membrane cells (Ctrl) where *ss* is repressed, with a median compaction of 588 nm (lower quartile = 456 nm; upper quartile = 720 nm). The *ss* locus was significantly more open in antennal cells (A) where *ss* is expressed, with a median compaction of 809 nm (lower quartile = 643 nm; upper quartile = 985 nm)(**Fig. 6D-E**). Thus, the DNA FISH method discerned differences in DNA compaction between cells with active or repressed *ss*.

### Chromatin compaction is dynamic during R7 differentiation

We examined compaction at the *ss* locus in the developing eye. In a single eye-antennal disc, we imaged all stages of R7 differentiation (**Fig. 2B**). As in previous experiments, we determined the differentiation state of cells based on their positions relative to the morphogenetic furrow. Undifferentiated cells were anterior to the morphogenetic furrow. Posterior to the morphogenetic furrow, precursors were located at 0-10 µm, differentiating cells were located at 10-30 µm, and R7s were located at >30 µm. R7s were also labeled by immunohistochemistry of a GFP reporter expressed in all R7s.

The *ss* locus was more compact in undifferentiated cells (*ss*^OFF^), similar to peripodial membrane cells (*ss*^OFF^) (**Fig. 6D**). The *ss* locus was more open in precursors (*ss*^ON^), similar to antennal cells (*ss*^ON^) (**Fig. 6D**). We predicted that differentiating cells (*ss*^OFF^) would be compact but were surprised to observe intermediate compaction (**Fig. 6D**). This intermediate compaction was also observed in R7s (a mix of *ss*^ON^ and *ss*^OFF^) (**Fig. 6D**). The intermediate compaction measurements suggested two main possibilities: (1) there are two distinct populations of cells, with compact or open chromatin at the *ss* locus, or (2) the *ss* locus is at an intermediate compaction state across all cells.

To discern between these hypotheses, we identified *ss*^ON^ and *ss*^OFF^ R7s using the reporter *Ins5* (**Fig. 4A**)*. ss*^ON^ R7s were identified based on *Ins5* expression. *ss*^OFF^ R7s were identified based on the absence of *Ins5* expression and their positions (see **Methods**). The *ss* locus was more open in *ss*^ON^ R7s (median = 786 nm), similar to other *ss*^ON^ cells, whereas the *ss* locus was more compact in *ss*^OFF^ R7s (550 nm), similar to other *ss*^OFF^ cells (**Fig. 6D-E**). These data suggested that the intermediate average compaction measurements observed for all R7s represented two distinct populations: *ss*^ON^ R7s with a more open *ss* locus and *ss*^OFF^ R7s with a more compacted *ss* locus.

Differentiating cells do not express *ss* and the variability in chromatin compaction prevented identification of two distinct cell populations. To characterize changes in compaction over time in differentiating cells, we examined “early” (at 10-20 µm) and “late” (at 20-30 µm) differentiating cells and observed no significant differences in compaction (**Fig. S6B-C**). Considering (1) the temporal progression of development from differentiating cells to R7s, and (2) the similarity of intermediate compaction between *ss*^OFF^ differentiating cells and the total population of R7s (including *ss*^ON^ and *ss*^OFF^ R7s), we surmise that differentiating cells also represented two populations: cells with a more open *ss* locus and cells with a more compacted *ss* locus. We cannot rule out that differentiating cells are comprised of cells with intermediate compaction states or a mix of cells with open, compact, and intermediate compaction states.

Comparing the expression and compaction states for developing R7s over time, we find that the *ss* locus is: (1) inactive and compact in undifferentiated cells, (2) active and open in precursors, (3) inactive and likely a mix of open and compact in differentiating cells, and (4) active and open, or inactive and compact in R7s. We next examined the relationship between transcription and chromatin compaction during R7 subtype specification.

### Transcription in precursors is required for decompaction of the *ss* locus

The early expression of *ss* in precursors driven by the *early enhancer* is required for expression later in R7s driven by the *late enhancer*. As no discernible Ss protein is generated during the early expression in precursors and Ss does not feedback to regulate its expression (**Fig. 3X-Z**)(Johnston and Desplan, 2014), we hypothesized that early expression in precursors was required to open the *ss* locus. To test this hypothesis, we examined promoter deletion mutants (*P*Δ) that do not express *ss* (**Fig. 3E-H, U-W**). In *P*Δ mutants, the *ss* locus was compact in undifferentiated cells and remained compact in precursors, differentiating cells, and R7s (**Fig. 6F**), suggesting that transcription plays a role in opening the *ss* locus in precursors. In *P*Δ mutants, the *ss* locus was similarly compact in antennal cells which normally express *ss* and have open chromatin (**Fig. 6F**). These data suggest that *ss* transcription is required for decompaction of the *ss* locus.

To test whether expression driven by the *early enhancer* is required to open the *ss* locus, we examined chromatin compaction in *EE*Δ mutants. In *EE*Δ mutants, *ss* was not expressed in precursors and R7s but remained active in the antenna (**Fig. 3I-L, U-W**). In *EE*Δ mutants, the *ss* locus was compact in undifferentiated cells, precursors, differentiating cells, and R7s but was open in antennal cells (**Fig. 6G**). Thus, early expression of *ss* in precursors driven by the *early enhancer* is required for decompaction of the *ss* locus in precursors, differentiating cells, and R7s.

We next tested the interaction of chromatin compaction and expression driven by the *late enhancer.* In *LE*Δ mutants, *ss* was expressed in precursors, turned off in differentiating cells, and remained off in all R7s (**Fig. 3M-P, U-W**). We hypothesized that either (1) the *ss* locus would display intermediate compaction in R7s, similar to differentiating cells, or (2) the *ss* locus would be more compacted similar to other *ss^OFF^* cells. In *LE*Δ mutants, the *ss* locus was compact in undifferentiated cells, open in precursors, and intermediate in differentiating cells and R7s (**Fig. 6H**), similar to wild type flies (**Fig. 6D**), suggesting that chromatin state in maturing R7s is not dependent on transcription driven by the *late enhancer*.

These data suggest that expression in precursors driven by the *early enhancer* is required to open the *ss* locus and that compaction state is independent of expression driven by the *late enhancer*.

## Discussion

### Temporally dynamic antagonism stochastically specifies R7 subtypes

We investigated how regulation of transcription and chromatin compaction at the *ss* locus controls stochastic R7 photoreceptor patterning in the fly eye. *ss* is initially in a compact, repressed state in undifferentiated cells in the developing eye. This compacted state is similar in other Ss^OFF^ cell types including peripodial cells and Ss^OFF^ R7s. As the eye develops, *ss* is transcribed in precursors and chromatin is opened. *ss* transcription and decompaction are lost in mutants deleting the *early enhancer* or the *promoter*, suggesting that *ss* transcription drives the opening of the *ss* locus in precursors.

As the cells mature, *ss* expression ceases and the *ss* locus compacts during the transition from the precursor to the differentiating cell phase. Our observations that (1) the *ss* locus is open in Ss^ON^ R7s and compact in Ss^OFF^ R7s and (2) the similarity of median compaction in differentiating cells and all R7s (including Ss^ON^ R7s and Ss^OFF^ R7s), suggest that the *ss* locus assumes either an open or compact state in differentiating cells that is maintained until terminal R7 subtype specification. Our data are consistent with stable compaction states in differentiating cells, but they cannot rule out changes during this phase of R7 development.

*ss* expression and compaction during the transition from precursor to differentiating cell phases are critical processes that determine the stochastic R7 fate choice. Decreasing *early enhancer* activity reduced *ss* expression in precursors and the proportion of Ss^ON^ R7s. Extending *ss* transcription into the differentiating cell phase increased the proportion of Ss^ON^ R7s. We propose that variable activation and duration of transcription in each precursor determines the probability of recompaction, which ultimately dictates the Ss^ON^ or Ss^OFF^ expression state in R7s.

In the last stage of R7 subtype specification, *ss* expression driven by the *late enhancer* is repressed in a subset of R7s. The repression reporter strategy showed that repression at the *ss* locus limits expression to a subset of Ss^ON^ R7s. The chromatin compaction assays showed that the *ss* locus is open in Ss^ON^ R7s and compact in Ss^OFF^ R7s. Together, our data suggest that chromatin compaction represses *ss* expression.

Based on these findings, we propose a mechanism that controls stochastic R7 subtype specification (**Fig. 7**). The *ss* locus is in a compact state in undifferentiated cells (**Fig. 7,** U). Transcription driven by the *early enhancer* opens the *ss* locus in precursors (**Fig. 7,** P). Early expression ceases and the *ss* locus randomly assumes an open or compact state in differentiating cells (**Fig. 7,** D). R7s with open chromatin at the *ss* locus reactivate *ss* and take on the Ss^ON^ R7 fate, whereas R7s with compact chromatin at the *ss* locus repress *ss* and take on the Ss^OFF^ R7 fate (**Fig. 7,** R7).

**Figure 7.**
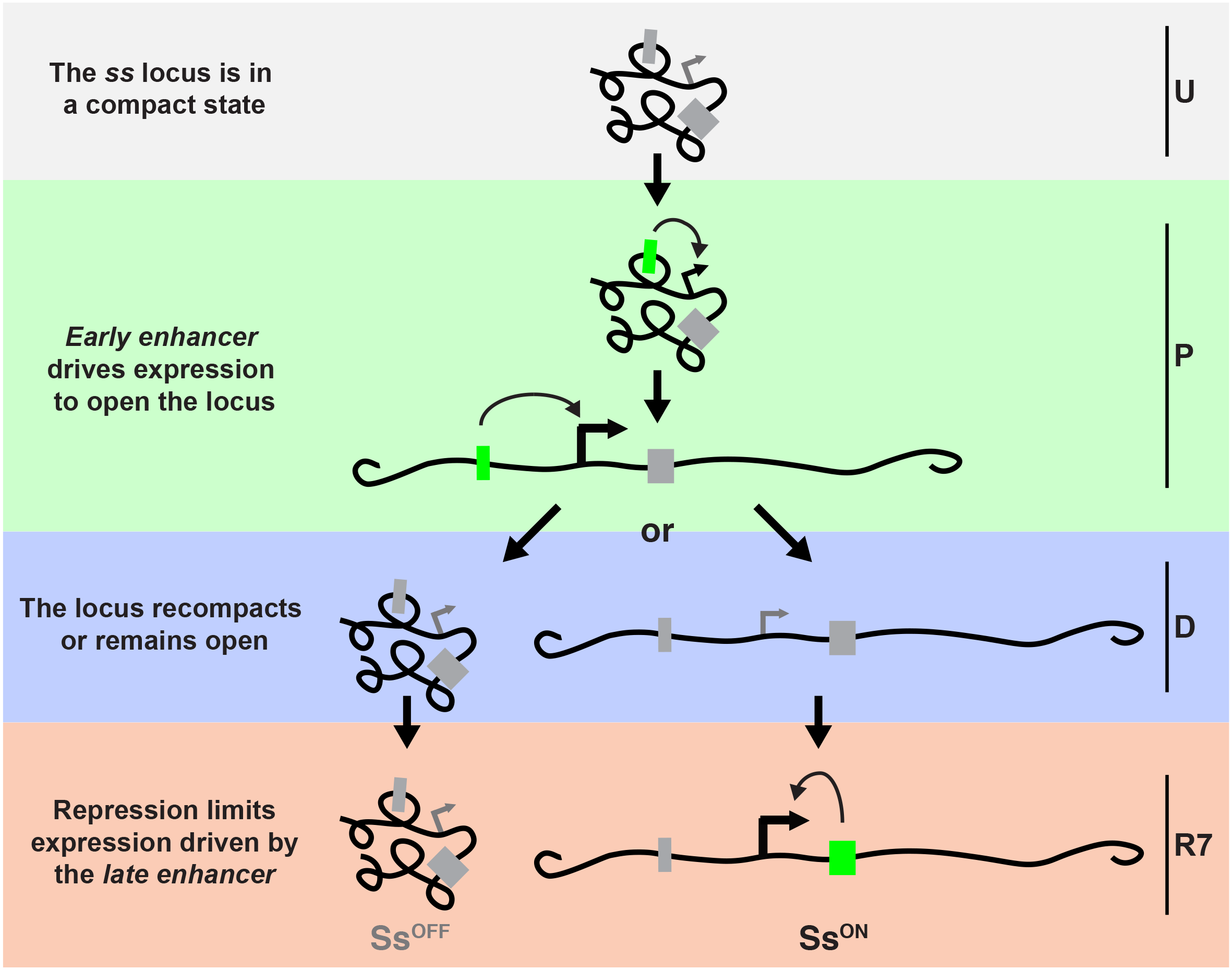
Proposed mechanism for stochastic R7 subtype specification. U = undifferentiated cells; P = precursors; D = differentiating cells; R7 = R7s. Gray box = inactive enhancer; green box = active enhancer.

### Prime and boost mechanisms controlling cell fate specification

A key aspect of this mechanism is the initial “priming” or opening of the *ss* locus during the early expression in precursors. Transcription-based priming plays important roles in several stereotyped developmental programs (Anderson et al., 2016; Cochella and Hobert, 2012; Greenberg et al., 2017; Kaikkonen et al., 2013; Schmitt et al., 2005). A well-understood example has been described in *C. elegans*, where the bilateral pair of ASE gustatory neurons display asymmetric gene expression and function (Ortiz et al., 2006, 2009). Stereotyped specification of the left neuron ASEL is dependent upon the asymmetric expression of the microRNA *lsy-6* (Johnston and Hobert, 2003), achieved by a “prime and boost” mechanism. Several cell divisions prior to the birth of the terminal ASEL neuron, a pulse of *lsy-6* expression in the precursor cell promotes decompaction of the *lsy-6* locus. This decompacted state is maintained in the ASEL lineage throughout development, allowing for reactivation of *lsy-6* in the terminal ASEL neuron. In the ASER lineage that never experiences the early pulse of *lsy-6* expression, the locus remains in a repressed, compacted state, preventing later activation by transcription factors that are expressed in both ASE neurons (Charest et al., 2020; Cochella and Hobert, 2012). Thus, early transcription of a key regulator (*lsy-6*) promotes one cell fate (ASEL) by antagonizing chromatin-mediated repression important for the specification of the alternative fate (ASER).

The transcription-based prime and boost mechanism controlling ASEL/R sensory neuron specification in *C. elegans* has many similarities to the mechanism that we have identified for R7 subtype specification. In both systems, early expression of a key regulator in precursor cells opens a locus (prime) so that it can be reactivated later upon terminal specification (boost). A major difference is that the ASEL/R decision requires priming in only the ASEL lineage to reproducibly generate the ASEL fate, whereas the R7 subtype decision utilizes priming in all precursors, which opens the chromatin followed by variable chromatin compaction and repression that ultimately determines the Ss^ON^ or Ss^OFF^ R7 fate.

Both the ASEL/R and R7 subtype decisions also exhibit a window of inactivity between the early and late expression phases. However, this window appears to play two very different roles. In the ASEL/R decision in worms, the early priming of the *lsy-6* locus occurs several cell divisions prior to terminal differentiation. The time between the prime and boost is an obstacle that must be overcome to remember the early developmental event. In contrast, the window between the early and late stages of *ss* expression appears to enable chromatin compaction and repression that determine the Ss^ON^ or Ss^OFF^ expression states in R7s.

### Shared features of stochastic fate specification

Though stochastic fate specification is an important feature of many cell fate programs, general features of these mechanisms have not been identified. In the bacterium *Bacillus subtilus*, transcriptional regulation is critical, as ComK transcription drives a stochastic cell fate switch to the “competent” fate. In both the competence decision in bacteria and R7 subtype specification in flies, all “precursor” cells express the key regulator, yet only a subset undergo the cell fate switch (Maamar et al., 2007; Mugler et al., 2016; Süel et al., 2006).

Stochastic R7 subtype specification in flies also shares mechanistic features with olfactory receptor selection in mice, particularly in the repression of alternative fates. In the olfactory system, OR genes are found in a compact heterochromatic region in the nucleus, with one gene that escapes repression and activates (Clowney et al., 2011; Magklara et al., 2011). Similarly, chromatin compaction and repression play key roles in determining *ss*^ON^ and *ss*^OFF^ R7 fates. Our studies in flies bridge the roles of transcription in bacteria and chromatin in mice for stochastic cell fate specification.

### Stochasticity and the antagonism between transcription and chromatin

Our understanding of the relationship between transcription and chromatin is often a chicken and egg problem: it is unclear whether transcription state dictates chromatin state or vice versa. Here, we provide evidence that clearly identifies these cause-effect relationships and show how they change during development. The *early enhancer* drives transcription to open chromatin in precursors. In differentiating cells, the *early enhancer* ceases to function and transcription stops. Chromatin remains open or closes, marking the stochastic step. Finally, a *late enhancer* turns on in mature R7s. In cells where the locus is open, transcription reinitiates, while in cells where the locus is closed, transcription is repressed. Thus, initially, transcription state regulates chromatin state and later, chromatin state controls transcription state.

Our studies not only outline this simple mechanism, but also identify how the stochastic step is regulated. The stochastic step occurs as cells cease *ss* transcription in the precursor phase and assume the open or compact chromatin state in differentiating cells. Changing early transcription alters the probability of chromatin closing and ultimately, the proportion of photoreceptor subtypes. Our findings provide an important step in understanding how transcription and chromatin states regulate one another to control how cells randomly assume fates.

## Materials and Methods

### RESOURCE AVAILABILITY

#### Lead Contact

All information queries or requests for resources can be directed to the Lead Contact, Robert Johnston Jr.

#### Materials Availability

All reagents and fly lines are available upon request.

#### Data and Code Availability

The script used to generate 19-bp bar-coding primers for Oligopaints probe design is available at https://github.com/kviets0913/Oligopaints-Primers-Custom-Script. The custom script used to analyze confocal images and determine the density of nascent RNA expression is available at https://github.com/lvoortman/Automated_Image_Analysis.

### KEY RESOURCE TABLE

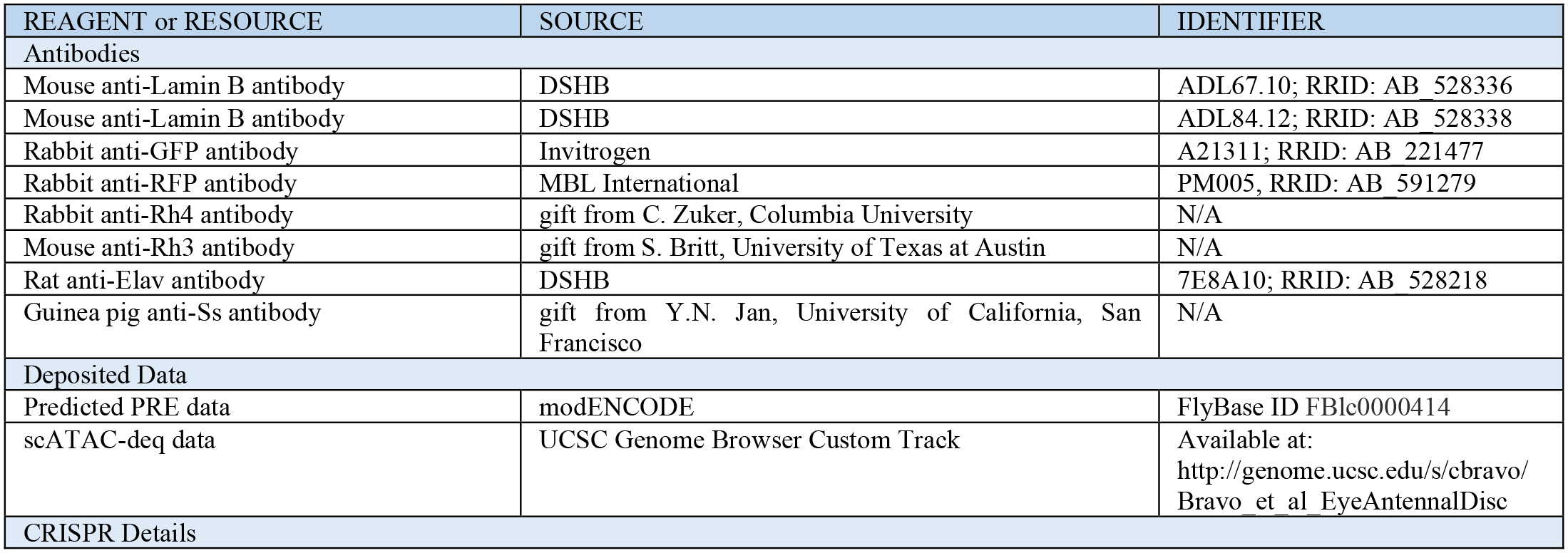
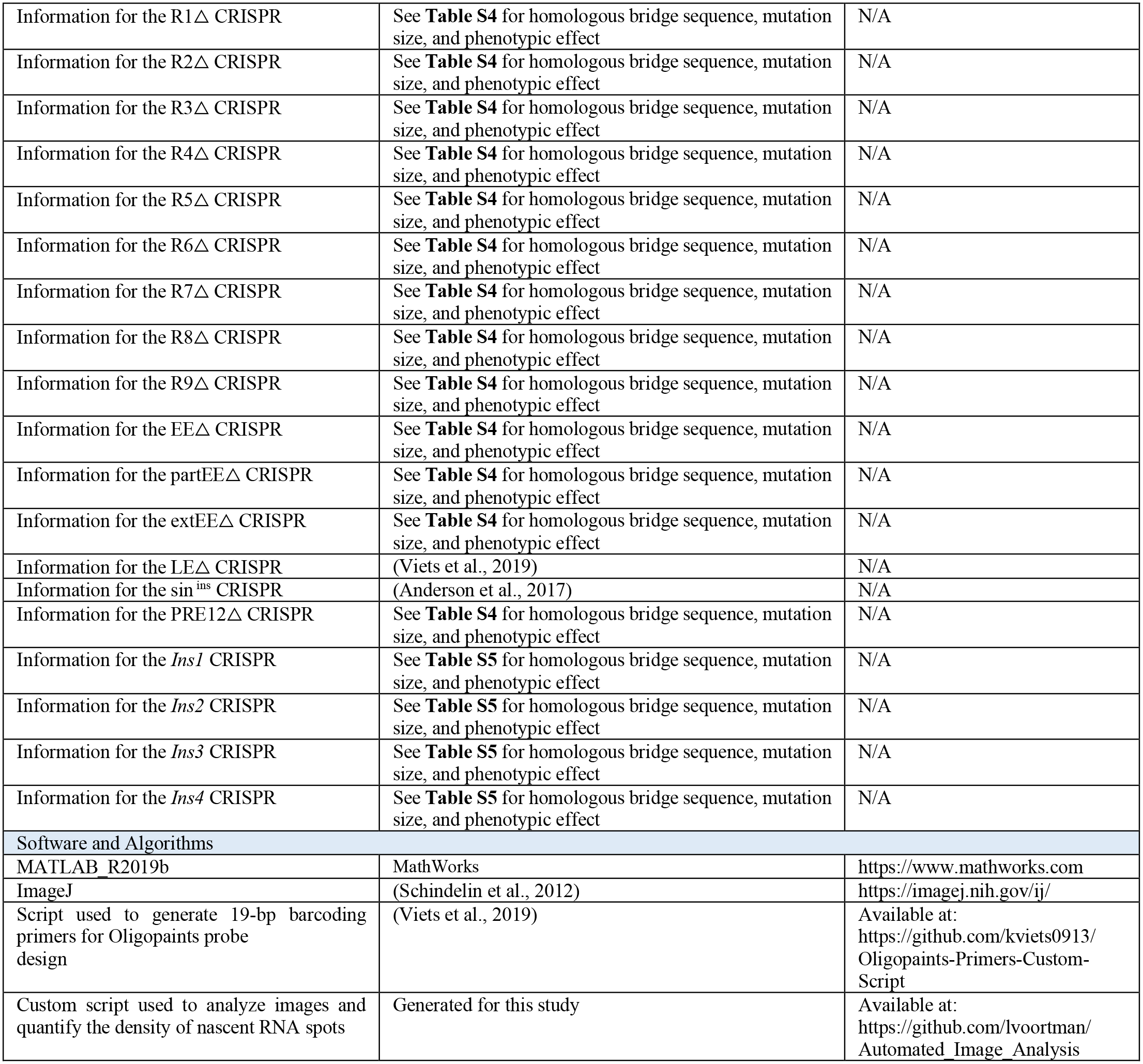

### EXPERIMENTAL MODEL AND SUBJECT DETAILS

#### Drosophila Lines

Flies were raised on standard cornmeal-molasses-agar medium and grown at 25° C. All experiments in this study included both male and female flies. See **Table S3** for a full list of fly genotypes used.

### METHOD DETAILS

#### Confocal Image Acquisition

All images were acquired using a Zeiss LSM 700 or LSM 980 confocal microscope. Adult retina images were acquired at a single Z plane at 20x magnification. Immunohistochemistry images at the pupal and larval stages were taken at 40x magnification as minimal Z stacks with a slice thickness of 500 nm. Image acquisition for larval DNA and RNA FISH experiments were taken at 63x magnification as large Z stacks encompassing the tissue with a slice thickness of 300nm.

#### CRISPR Mediated Deletions

Deletions to the endogenous *ss* locus were generated using CRISPR (Gratz et al., 2014; Port et al., 2014). Sense and antisense DNA Information for the forward and reverse strands of the gRNA were designed to generate BbsI restriction site overhangs. The oligos were annealed and cloned into the pCFD3 cloning vector (Addgene, Cambridge, MA). A single stranded DNA homology bridge was generated with 60 bp homologous regions flanking each side of the predicted cleavage site. The gRNA construct (500 ng/µl) and homology bridge oligo (100 ng/µl) were injected into Drosophila embryos (BestGene, Inc.). Single males were crossed with a balancer stock (yw; +; TM2/TM6B), and F1 female progeny were screened for the deletion via PCR and sequencing. Single F1 males whose siblings were deletion-positive were crossed to the balancer stock (yw; +; TM2/TM6B) and the F2 progeny were screened for the deletion via PCR and sequencing. Information on all CRISPR oligonucleotides used for this study can be found in **Table S4**.

#### CRISPR Mediated Insertions

Insertion into the endogenous *ss* locus were generated using CRISPR. sgRNAs were designed using Chopchop and cctop (Labuhn et al., 2018; Labun et al., 2016), isolated from injection stocks, and amplified using in vitro transcription. A single stranded DNA homology bridge was generated with homologous regions flanking each side of the predicted cleavage site after inserted into the pBluescript SK vector backbone using Gibson Assembly. The homology bridge oligos were co-injected with Cas9 RNA (2 µg/µl) and sgRNA (1 µg/µl) into 300 Drosophila embryos (Qidong Fungene Biotechnology Co., Ltd.). Single males were crossed with a balancer stock (yw; +; TM2/TM6B), and F1 female progeny were screened for the deletion via PCR and sequencing. Single F1 males whose siblings were deletion-positive were crossed to the balancer stock (yw; +; TM2/TM6B) and the F2 progeny were screened for the deletion via PCR and sequencing. Information on all CRISPR oligonucleotides used for this study can be found in **Table S5**.

#### Antibodies

Antibodies and dilutions were as follows: mouse anti-lamin B (DSHB ADL67.10 and ADL84.12), 1:100; rabbit anti-GFP (Invitrogen), 1:500; rabbit anti-RFP (MBL), 1:400 rat anti-Elav (DSHB, 7E8A10), 1:50; rabbit anti-Rh4 (gift from C. Zuker, Columbia University), 1:50; mouse anti-Rh3 (gift from S. Britt, University of Texas at Austin), 1:50; guinea pig anti-Ss (gift from Y.N. Jan, University of California, San Francisco), 1:500; all secondary antibodies (Molecular Probes) were Alexa Fluor-conjugated and used at 1:400.

#### Immunohistochemistry

Adult, mid-pupal, and larval retinas were dissected as described (Hsiao et al., 2012) and fixed for 15 min with 4% formaldehyde at room temperature. Retinas were rinsed three times in PBS plus 0.3% Triton X-100 (PBX) followed by three 15 min washes in PBX. Retinas were incubated with primary antibodies diluted in PBX >2hrs at room temperature and then rinsed three times in PBX followed by three 15 min washes in PBX. Retinas were incubated with secondary antibodies diluted in PBX >2hrs at room temperature and then rinsed three times in PBX followed by three 15 min washes in PBX. Retinas were mounted in SlowFade Gold Antifade Reagent (Invitrogen). Images were acquired using a Zeiss LSM 700 or LSM 980 confocal microscope at 20x or 40x magnification.

#### Enhancer Reporter

The *early enhancer* and *late enhancer* cassettes were amplified and inserted into the pJR20 plasmid. The *early enhancer* was amplified from chr3R:16,410,464 - 16,411,045. The *late enhancer* was amplified from chr3R:16,399,856-16,396,676. All plasmids used were made through standard cloning procedures. Plasmids, plasmid maps, and cloning details are available on request. All constructs were sent to BestGene (Chino Hills, CA) for injection. Constructs were inserted via PhiC31 integration at the attP40 landing site.

#### Oligopaints Probe Design for RNA and DNA FISH

Probes for RNA and DNA FISH were designed using the Oligopaints technique (Beliveau et al., 2012, 2013, 2015). Target sequences were run through the bioinformatics pipeline available at http://genetics.med.harvard.edu/oligopaints/ to identify sets of 50-bp optimized probe sequences (i.e. ‘‘libraries’’) tiled across the DNA sequence of interest. Five 19-bp bar-coding primers, gene F and R; universal (univ) F and R, and either sublibrary (sub) F or random (rando) R, were appended to the 5’ and 3’ ends of each probe sequence. To ensure that all probes were the same length, an additional 8-bp random sequence was added to the 3’ end of the probes. The gene F and R primers allowed PCR amplification of a probe library of interest out of the total oligo pool, and the univ F and R primers allowed conjugation of fluorophores, generation of single-stranded DNA probes, and PCR addition of secondary sequences to amplify probe signal. The ss 50-kb left and right extension libraries had a sub F primer between the gene and universal forward primers to allow PCR amplification of probes targeting a specific sub-region of the locus of interest. All other probe libraries had a rando R primer appended at the 3’ end to maintain a constant sequence length between all probes. Bar-coding primer sequences were taken from a set of 240,000 randomly generated, orthogonal 25-bp sequences (Xu et al., 2009) and run through a custom script (available at https://github.com/kviets0913/Oligopaints-Primers-Custom-Script) to select 19-bp sequences with 15-bp homology to the Drosophila genome. Primers were appended to probe sequences using the orderFile.py script available at http://genetics.med.harvard.edu/oligopaints/. Completed probe libraries were synthesized as custom oligo pools by Custom Array (Bothell, WA), and fluorescent FISH probes were generated as described in references (Beliveau et al., 2012, 2013, 2015; Viets et al., 2019).

#### RNA FISH

RNA FISH was performed using modified versions of the protocols described in references (Beliveau et al., 2012, 2015). 20-50 eye/antennal discs attached to mouth hooks from third instar larvae were collected on ice and fixed in 129 µL ultrapure water, 20 µL 10X PBS, 1 µL Tergitol NP-40, 600 µL heptane, and 50 µL fresh 16% formaldehyde. Tubes containing the fixative and eye discs were shaken vigorously by hand, then fixed for 10 minutes at room temperature with nutation. Eye discs were then given three quick washes in 1X PBX, followed by three five-minute washes in PBX with 0.5% (vol/vol) RNAse inhibitor (Promega) at room temperature with nutation. Eye discs were then removed from the mouth hooks and blocked for 1 hour in 1X PBX:Western Blocking Reagent (Roche) at room temperature with nutation. They were then incubated in primary antibody diluted in 1X PBX with 0.5 U/pL RNAse inhibitor overnight at 4° C with nutation. Next, eye discs were washed three times in 1X PBX for 20 minutes and incubated in secondary antibody diluted in 1X PBX with 0.5 U/pL RNAse inhibitor for two hours at room temperature with nutation. Eye discs were then washed two times for 20 minutes in 1X PBX, followed by a 20-minute wash in 1X PBS. Next, discs were given one 10-minute wash in 20% formamide + 80% 2X SSCT (2X SSC+.001% Tween-20), one 10-minute wash in 40% formamide + 60% 2X SSCT, and two 10-minute washes in 50% formamide + 50% 2X SSCT. Discs were then predenatured by incubating for four hours at 37° C, three minutes at 92° C, and 20 minutes at 60° C. Primary probes were added in 36 µL hybridization buffer consisting of 50% formamide + 50% 2X SSCT+2% dextran sulfate (w/v). All probes were added at a concentration of ≥5 pmol fluorophore/mL. 4 µL of probe was added. After addition of probes, eye discs were incubated at 37° C for 16-20 hours with shaking. Eye discs were then washed for 1 hour at 37° C with shaking in 50% formamide + 50% 2X SSCT. 1 µL of each secondary probe was added at a concentration of 100 pmol/mL in 50 µL of 50% formamide + 50% 2X SSCT. Secondary probes were hybridized for 1 hour at 37° C with shaking. Eye discs were then washed twice for 30 minutes in 50% formamide + 50% 2X SSCT at 37° C with shaking, followed by three 10-minute washes at room temperature in 20% formamide + 80% 2X SSCT and 2X SSCT with nutation. Discs were incubated in 2X SSCT with 300 µM DAPI for 15 minutes at room temperature with nutation, followed by three 10-minute washes at room temperature in 2X SSC with nutation. Discs were mounted in SlowFade Gold immediately after the final 2X SSC wash and imaged using a Zeiss LSM700 or Zeiss LSM980 confocal microscope at 63x magnification.

#### DNA FISH

DNA FISH was performed using modified versions of the protocols described in references (Beliveau et al., 2012, 2013, 2015; Viets et al., 2019). 20-50 eye/antennal discs attached to mouth hooks from third instar larvae were collected on ice and fixed in 129 µL ultrapure water, 20 µL 10X PBS, 1 µL Tergitol NP-40, 600 µL heptane, and 50 µL fresh 16% formaldehyde. Tubes containing the fixative and eye discs were shaken vigorously by hand, then fixed for 10 minutes at room temperature with nutation. Eye discs were then given three quick washes in 1X PBX, followed by three five-minute washes in PBX at room temperature with nutation. Eye discs were then removed from the mouth hooks and blocked for 1 hour in 1X PBX+1% BSA at room temperature with nutation. They were then incubated in primary antibody diluted in 1X PBX overnight at 4° C with nutation. Next, eye discs were washed three times in 1X PBX for 20 minutes and incubated in secondary antibody diluted in 1X PBX for two hours at room temperature with nutation. Eye discs were then washed two times for 20 minutes in 1X PBX, followed by a 20-minute wash in 1X PBS. Next, discs were given one 10-minute wash in 20% formamide + 80% 2X SSCT (2X SSC+.001% Tween-20), one 10-minute wash in 40% formamide + 60% 2X SSCT, and two 10-minute washes in 50% formamide + 50% 2X SSCT. Discs were then predenatured by incubating for four hours at 37° C, three minutes at 92° C, and 20 minutes at 60° C. Primary probes were added in 36 µL hybridization buffer consisting of 50% formamide + 50% 2X SSCT+2% dextran sulfate (w/v), + 1 µL RNAse A. All probes were added at a concentration of ≥5 pmol fluorophore/mL. For FISH experiments in which a single probe was used, 4 µL of probe was added. For FISH experiments in which three probes were used, 1.3 µL of each probe was added. After addition of probes, eye discs were incubated at 91° C for three minutes and at 37° C for 16-20 hours with shaking. Eye discs were then washed for 1 hour at 37° C with shaking in 50% formamide + 50% 2X SSCT. 1 µL of each secondary probe was added at a concentration of 100 pmol/mL in 50 mL of 50% formamide + 50% 2X SSCT. Secondary probes were hybridized for 1 hour at 37° C with shaking. Eye discs were then washed twice for 30 minutes in 50% formamide + 50% 2X SSCT at 37° C with shaking, followed by three 10-minute washes at room temperature in 20% formamide + 80% 2X SSCT and 2X SSCT with nutation. Discs were incubated in 2X SSCT with 300 µM DAPI for 15 minutes at room temperature with nutation, followed by three 10-minute washes at room temperature in 2X SSC with nutation. Discs were mounted in SlowFade Gold immediately after the final 2X SSC wash and imaged using a Zeiss LSM700 or Zeiss LSM980 confocal microscope at 63x magnification.

#### scATACseq analysis

Regions of the eye/antennal disc containing open chromatin were obtained from http://genome.ucsc.edu/s/cbravo/Bravo_et_al_EyeAntennalDisc (Bravo González-Blas et al., 2020) and viewed on the UCSC Genome Browser (WJ et al., 2002). Only regions of the eye/antennal imaginal disc containing *ss* expression (antennal cells, precursors, and mature photoreceptors) were compared. All tracks were scaled to the same parameters for accurate comparisons.

### QUANTIFICATION AND STATISTICAL ANALYSIS

#### Adult Eye Quantifications

The frequencies of Rh4- and Rh3-expressing R7s were scored manually for at least five eyes per genotype. R7s co-expressing Rh3 and Rh4 were scored as Rh4-positive (Mazzoni et al., 2008; Thanawala et al., 2013). 100 or more R7s were scored for each eye. The co-expression of reporters and Rh4- or Rh3-expressing R7s were scored manually for at least five adult eyes per genotype. 100 or more R7s were scored for each eye.

#### Density of Expression Quantification

The density of *ss* RNA punctae were calculated computationally using a custom written script in MATLAB. Due to homologous chromosome pairing between both copies of the endogenous chromosome, we observed a single dot for each chromosome. All images were acquired as 3D z-stacks with a slice thickness of 300 nm. The most highly expressed 25 slices within the antenna, and all slices containing punctae in R7 precursors were maximum intensity projected. Undifferentiated, precursor, and differentiating cells were demarcated in 10 µm regions based on distance from the morphogenetic furrow. The punctae were then identified and the area was determined as a bounding box encompassing the identified spots. Nascent RNA spots were distinguished from mature transcripts using an intensity threshold, removing them from density calculations. The density was calculated as the number of punctae per unit area in µm^2^. To ensure high fidelity identification of spots, four images were quantified manually in parallel, and the number of spots in each image were compared as the percentage of manual IDs. Automated identification had a mean %ID of 100.02% +/-0.67% when compared to manual quantification, indicating high fidelity identification and density quantification.

#### Compaction Quantification

All compaction quantifications were performed in 3D on z-stacks with a slice thickness of 300 nm. Quantifications were performed manually using Fiji (Joyce et al., 2012; Schindelin et al., 2012; Schneider et al., 2012). Boundaries were drawn for each image to denote cell type. Undifferentiated, precursor, and differentiating cells were demarcated in 10 µm regions based on distance from the morphogenetic furrow. To determine the z position of each FISH dot, an encapsulating box was drawn around the dot and the Plot Profile tool was used to assess the stack in which the dot was brightest. Due to homologous chromosome pairing between both copies of the endogenous chromosome, we observed a single dot for each DNA region. To determine the x-y-z distance between FISH dots, we used the multipoint tool to mark the center position for each spot within each nucleus. The distance between the FISH dots was then calculated in 3D as:

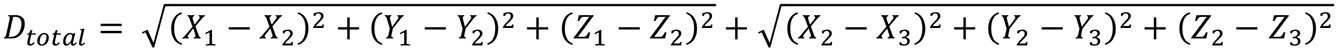

#### Statistical Analysis

All datasets were tested for a Gaussian distribution using a D’Agostino and Pearson omnibus normality test and a Shapiro-Wilk normality test. If either test indicated a non-Gaussian distribution for any of the datasets in an experiment, datasets were tested for statistical significance using a Wilcoxon rank-sum test (for single comparisons) or a one-way ANOVA on ranks with Dunn’s multiple comparisons test (for multiple comparisons). If both the D’Agostino and Pearson and the Shapiro-Wilk tests indicated a Gaussian distribution for all datasets in an experiment, datasets were tested for statistical significance using an unpaired t-test with Welch’s correction (for single comparisons) or an ordinary one-way ANOVA with Dunnett’s multiple comparisons test (for multiple comparisons). These statistical tests were performed using GraphPad Prism. Statistical tests and p-values are described in the figure legends.

## Supplemental Figure Legends

**Figure S1.**
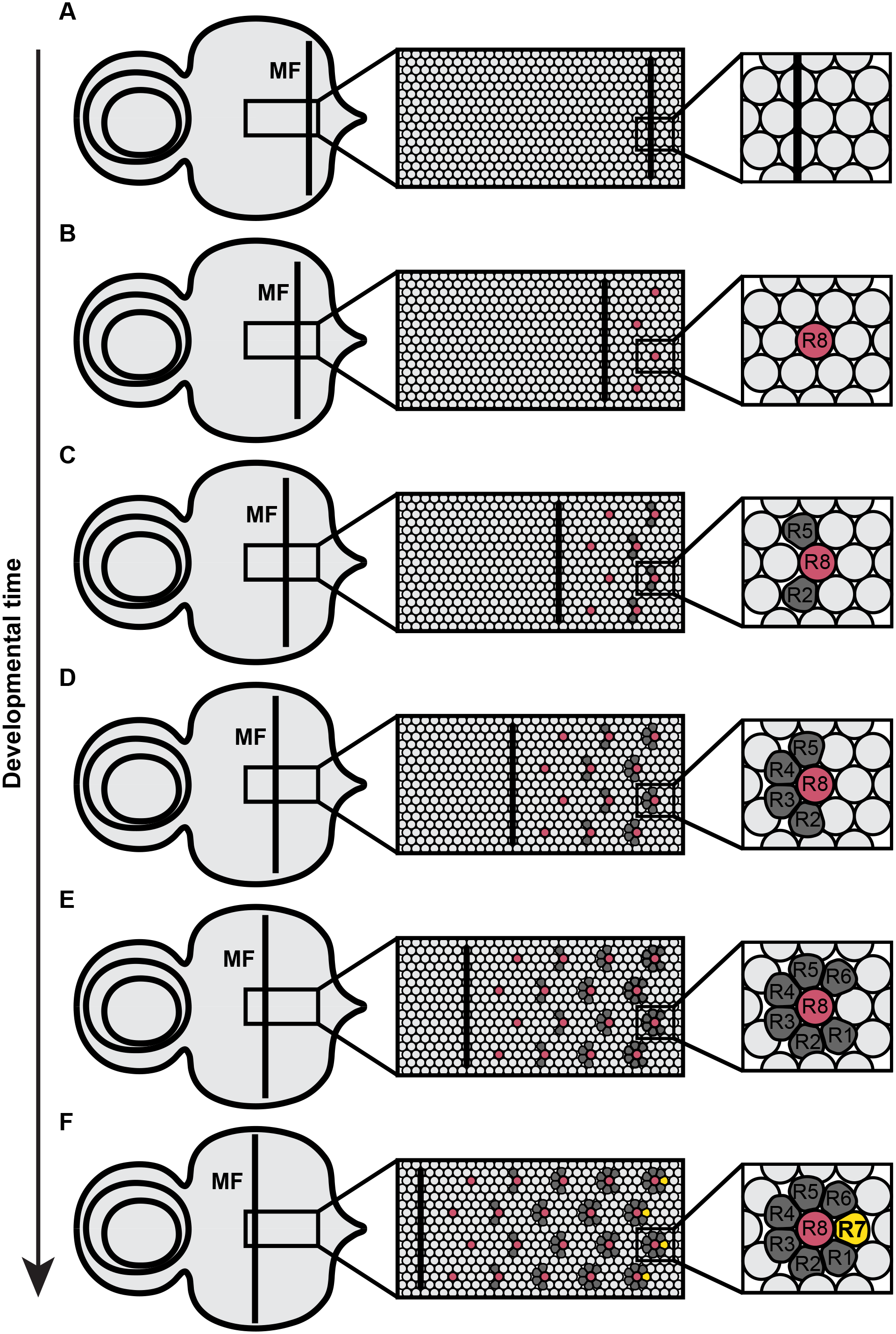
Schematic of photoreceptor specification in the eye/antennal disc. Related to Figure 1. **A-F)** Photoreceptor specification and morphogenetic furrow during eye development. Light gray circle = unspecified cell; maroon circle = R8 cell; dark gray circle = outer photoreceptor cell (R1-R6); yellow circle = R7 cell.

**Figure S2.**
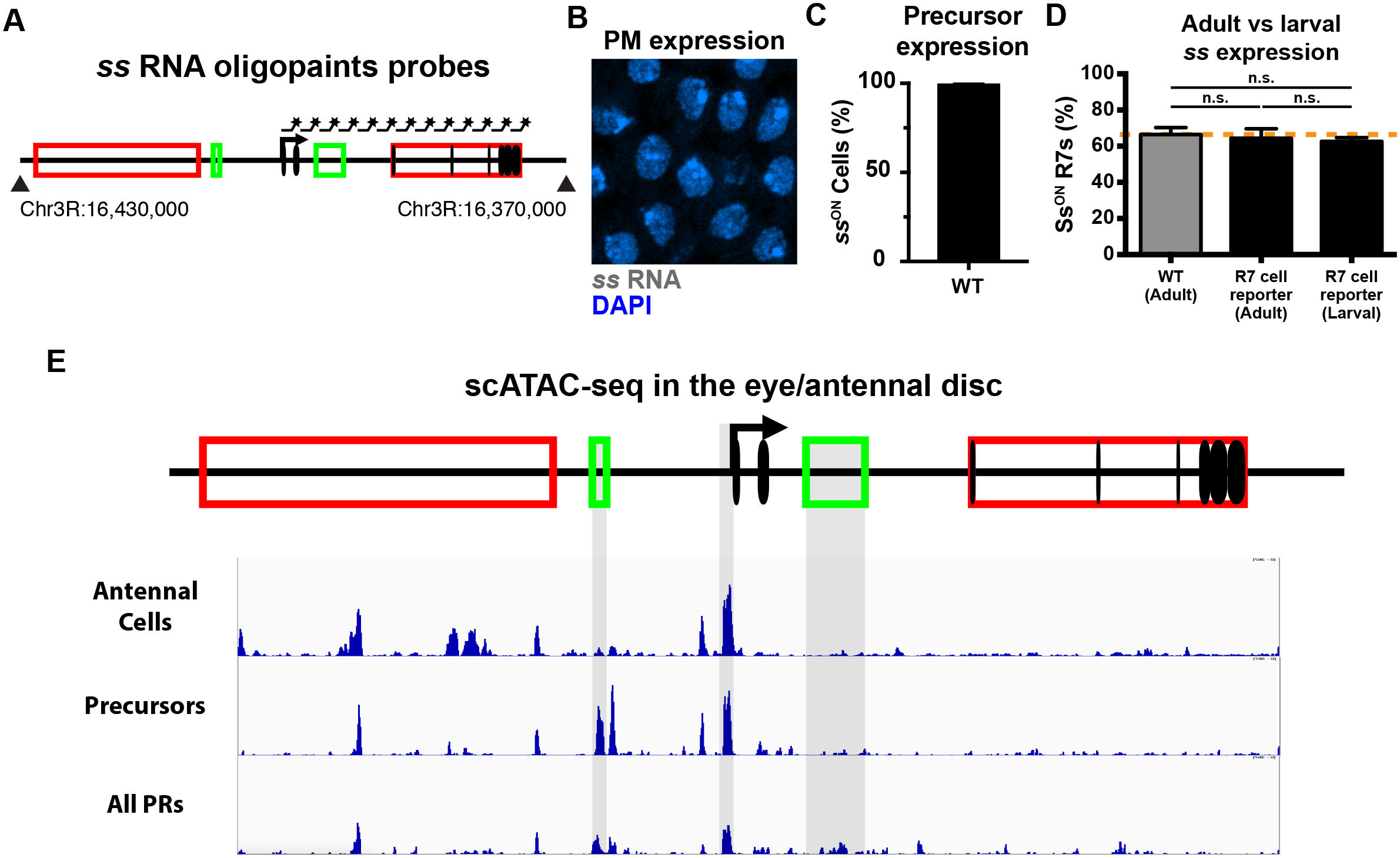
*ss* expression analysis. Related to Figure 2. **A)** Schematic of probes used to label *ss* RNA. **B)** *ss* is not expressed in peripodial membrane cells (*ss*^OFF^ control). Gray = *ss* RNA; blue = DAPI. **C)** All precursors transcribe *ss* in WT animals. **D)** % Ss^ON^ R7s is similar in adult R7s in WT flies, in adult R7s in the R7 cell reporter line, and in larval R7s in the R7 cell reporter line. Orange line indicates mean % Ss^ON^ R7s in adults. n.s. denotes p > 0.05. **E)** scATAC-seq reads at the *ss* locus by region of the eye/antennal disc. Gray rectangles = peaks matching *cis* regulatory elements. PRs = photoreceptors.

**Figure S3.**
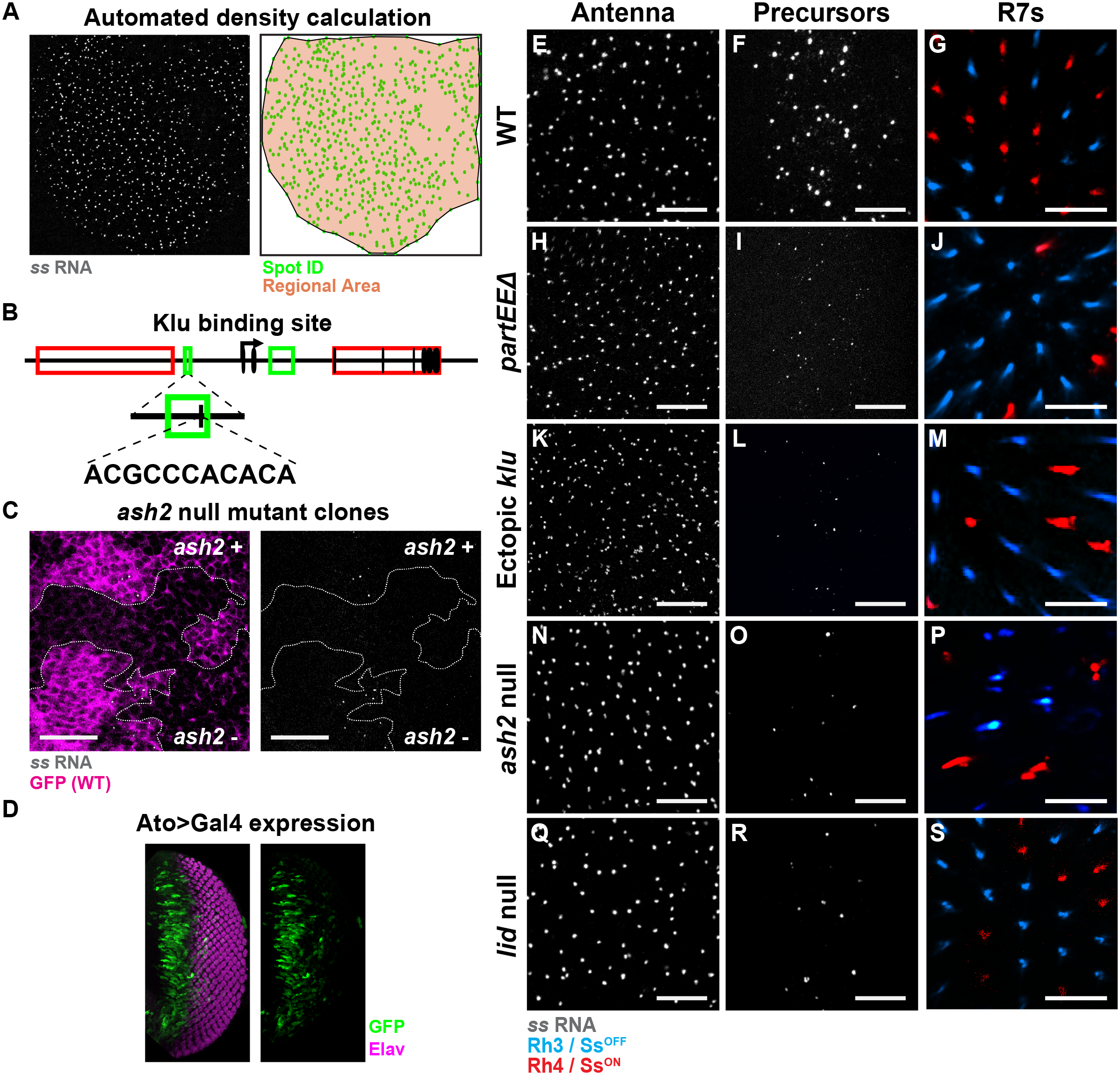
Automated density calculation; decreasing early *ss* expression decreases the proportion of Ss^ON^ R7s. Related to Figure 2. **A)** Automated identification and calculation of *ss* nascent RNA expression and density. **B)** Schematic of the Klu binding site and binding motif in the *early enhancer* of the *ss* locus. **C)** Early *ss* RNA expression is reduced in precursors in *ash2* null mutant clones. GFP-indicates *ash2* mutant clones; GFP+ indicates wild type clones. Magenta = GFP; dashed line = clone boundary. Gray = *ss* RNA; green = spot ID; orange = regional area. **D)** Ato>Gal4 drives GFP in precursors. Green = GFP; magenta = Elav. **For E, F, H, I, K, L, N, O, Q, R)** Scalebar = 10 µm. **For E, H, K, N, Q)** *ss* RNA expression in the antenna. Gray = *ss* RNA. **For F, I, L, O, R)** Early *ss* RNA expression in precursors. Gray = *ss* RNA. **For G, J, M, P, S)** Adult Rh3/Ss^OFF^ and Rh4/Ss^ON^ expression in R7s. Blue = Rh3; red = Rh4. **E-G)** In wild type (WT) animals, *ss* is expressed in antennal cells, precursors, and a subset of R7s. **H-J)** *pEED/def* mutants display normal *ss* expression in antennal cells, reduced *ss* expression in precursors, and a decrease in % Rh4/Ss^ON^ R7s. **K-M)** Animals with ectopic expression of Klu display normal *ss* expression in antennal cells, reduced *ss* expression in precursors, and a decrease in % Rh4/Ss^ON^ R7s. **N-P)** *ash2* null mutants display normal *ss* expression in antennal cells, reduced *ss* expression in precursors, and a decrease in % Rh4/Ss^ON^ R7s. **Q-S)** *lid* null mutant*s* display normal *ss* expression in antennal cells, reduced *ss* expression in precursors, and a decrease in % Rh4/Ss^ON^ R7s.

**Figure S4.**
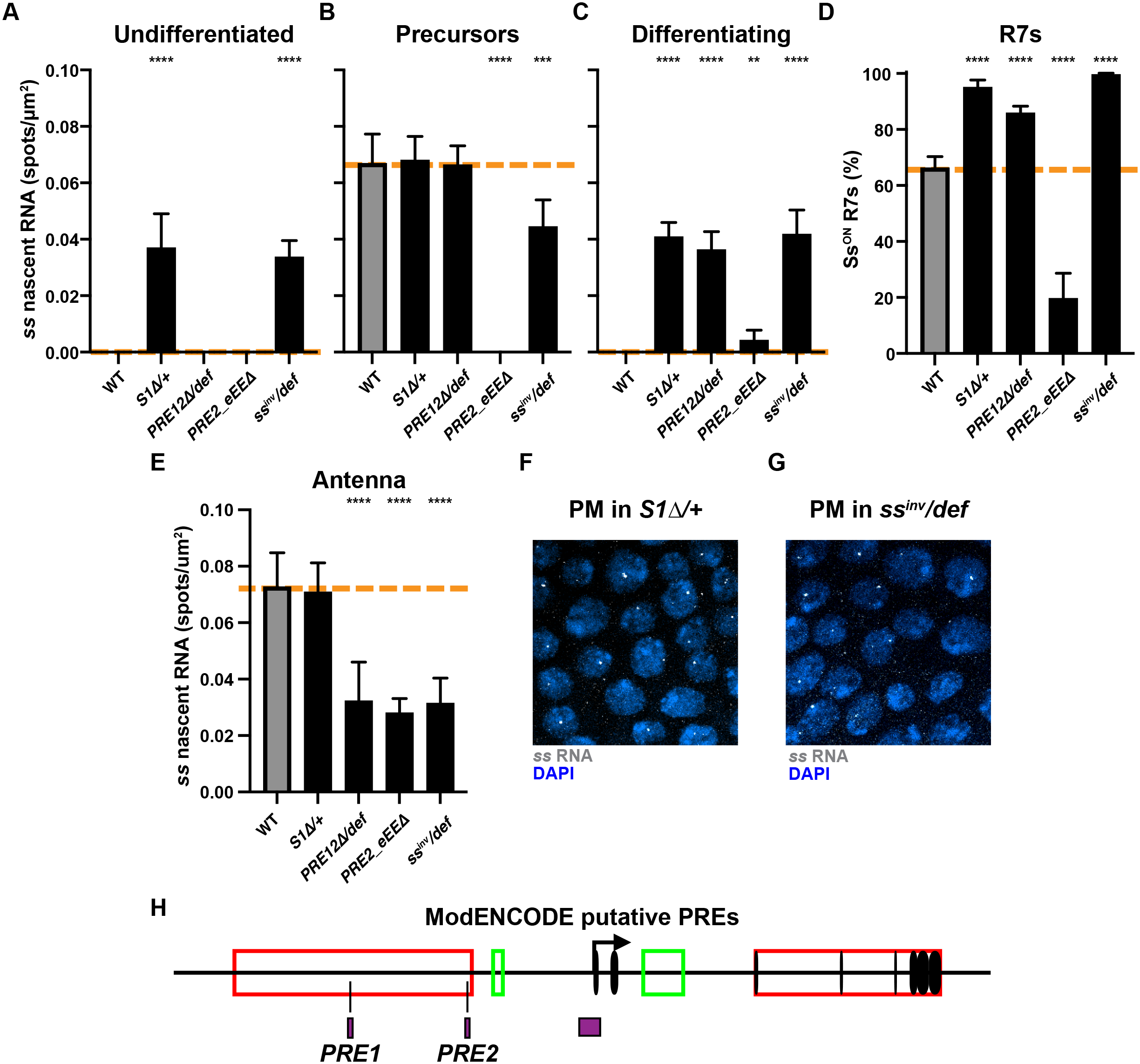
Derepressing early *ss* expression increases the proportion of Ss^ON^ R7s; putative PREs in the *ss* locus. Related to Figure 3. **For A-E)** Quantification of *ss* expression. Orange line indicates mean WT *ss* expression. ** denotes p < 0.005; *** denotes p < 0.0005; **** denotes p < 0.0001. **A)** Undifferentiated cells. **B)** Precursors **C)** Differentiating cells. **D)** R7s **E)** Antennal cells. **F-G)** *ss* is expressed in peripodial membrane cells in *S1D* and *ss^inv^* mutants. Gray = *ss* RNA; blue = DAPI. **H)** Putative PREs in the *ss* locus. Data adapted from FlyBase Jbrowse modENCODE plug-in. Purple rectangles indicate putative PREs.

**Figure S5.**
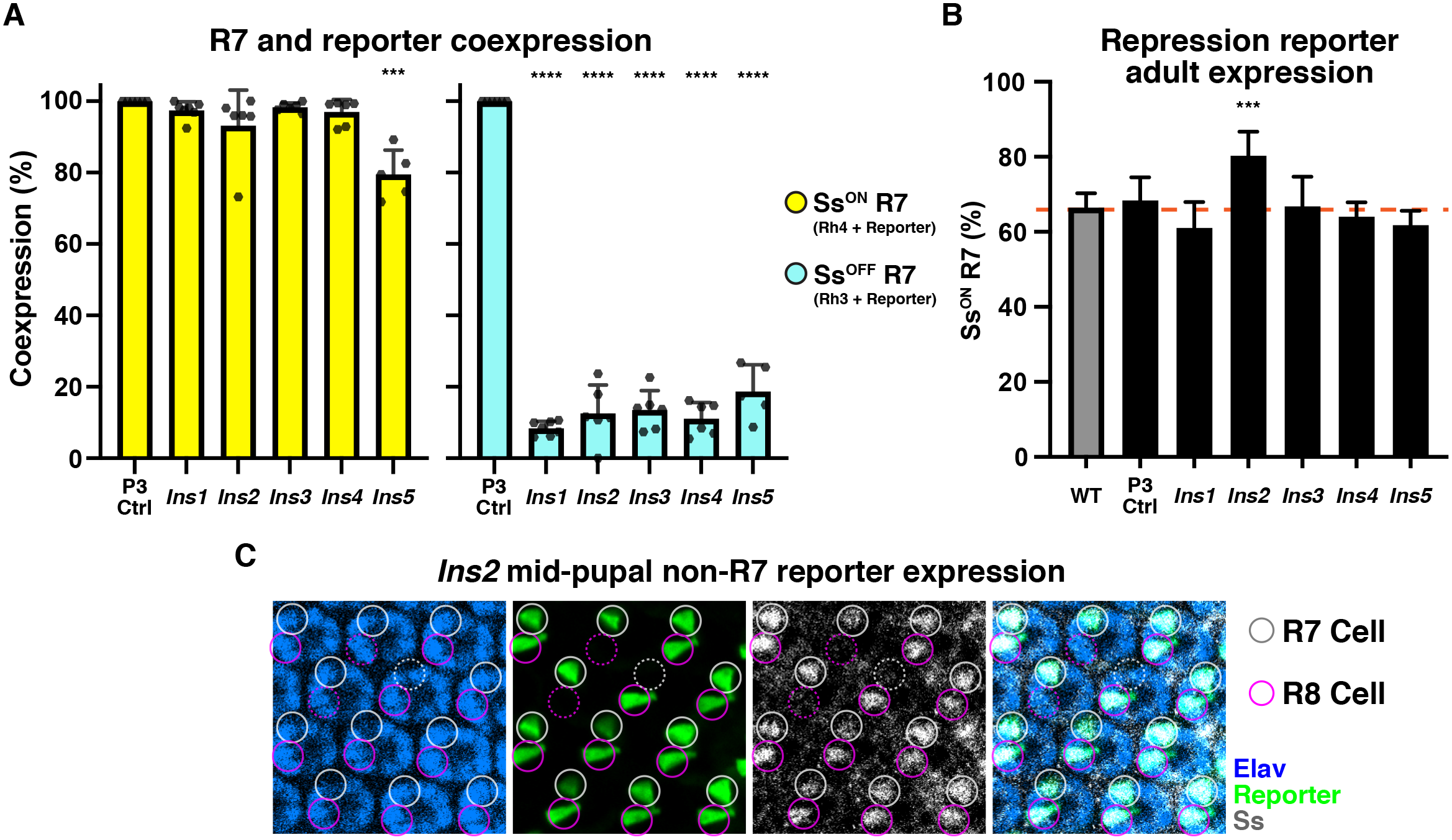
Repression by the *ss* locus limits expression to a subset of R7s; reporter and Ss are expressed in R8s in flies carrying *Ins2.* Related to Figure 4. **A)** Quantification of rhodopsin and reporter co-expression. Cyan = Rh3/Ss^OFF^, reporter^ON^; yellow = Rh4/Ss^ON^, reporter^ON^. **B)** % Rh4/Ss^ON^ R7s in control flies or flies carrying the repression reporter. Orange line indicates mean WT Ss expression. **C)** The reporter and Ss are expressed in R8s in flies carrying *Ins2* in mid-pupal retinas. Gray circles = R7s; maroon circles = R8s; blue = Elav; green = reporter; gray = Ss. Solid lines indicate reporter expressing cell; dotted lines indicate no reporter expression.

**Figure S5.**
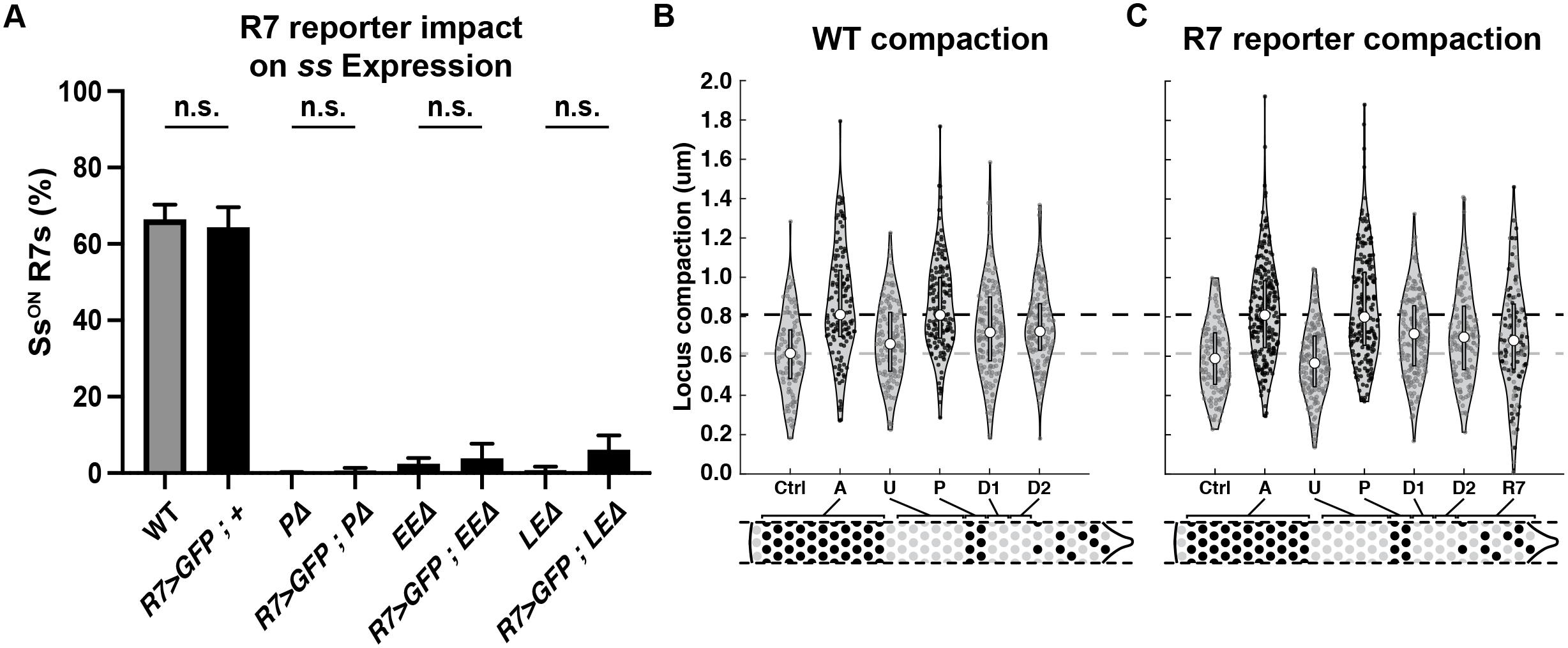
Control chromatin compaction and *ss* expression experiments; scATAC-seq analysis at the *ss* locus. Related to Figure 5. **A)** %Rh4/ Ss^ON^ R7s in WT and mutant genetic conditions with and without the R7 reporter. n.s. denotes p > 0.05. **B-C)** *ss* locus compaction dynamics are similar in WT and R7 reporter flies. Ctrl = peripodial membrane cells; A = antennal cells; U = undifferentiated cells; P = precursors; D = differentiating cells; R7 = R7s. Black circle = *ss*^ON^ cell; gray circle = *ss*^OFF^ cell; gray rectangle = quartile; white circle = median; gray dashed line = *ss*^OFF^ control/peripodial membrane median; black dashed line = *ss*^ON^ control/antennal cells median.

**Table S1.**
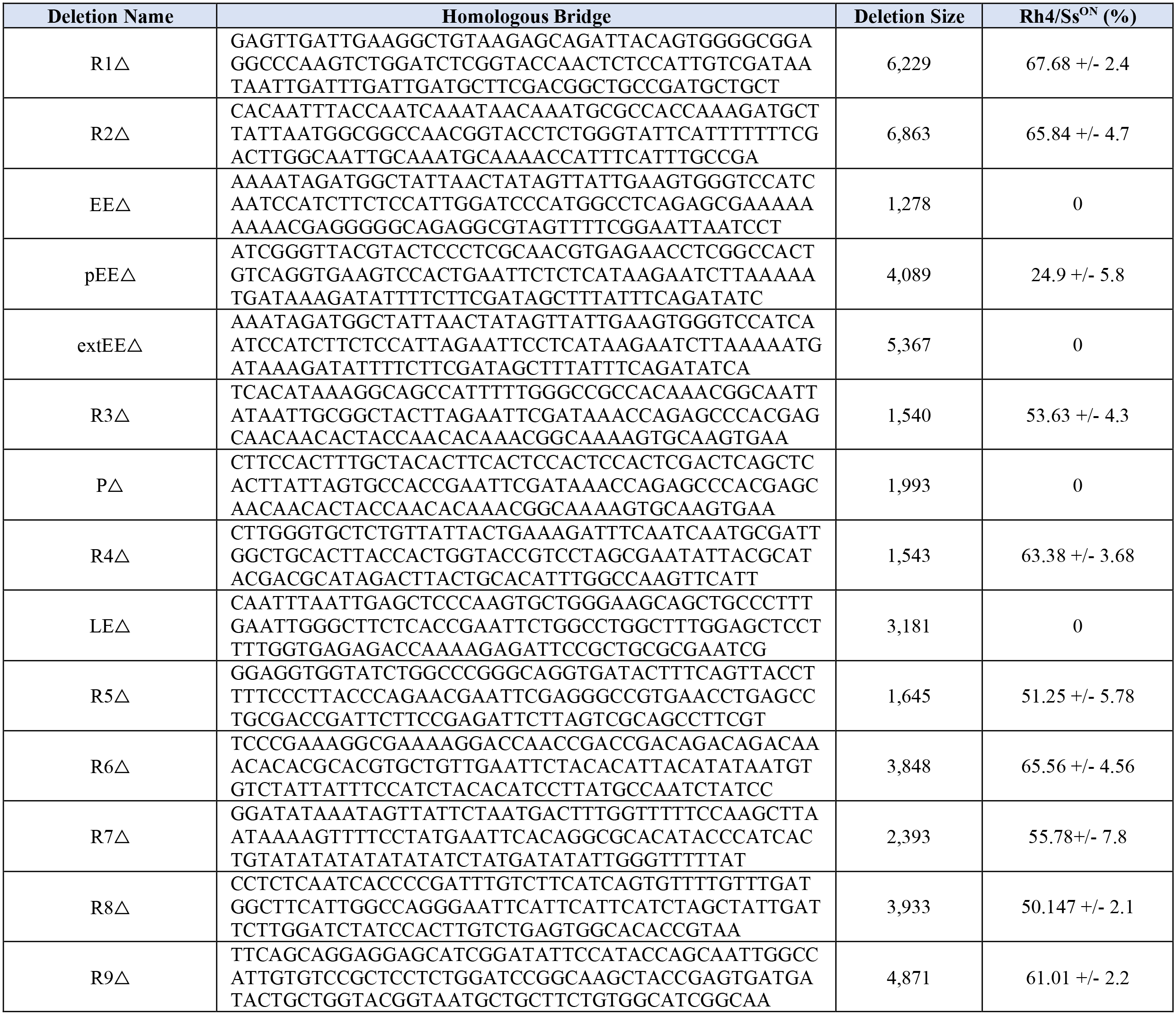
CRISPR deletion screen.

**Table S2.**
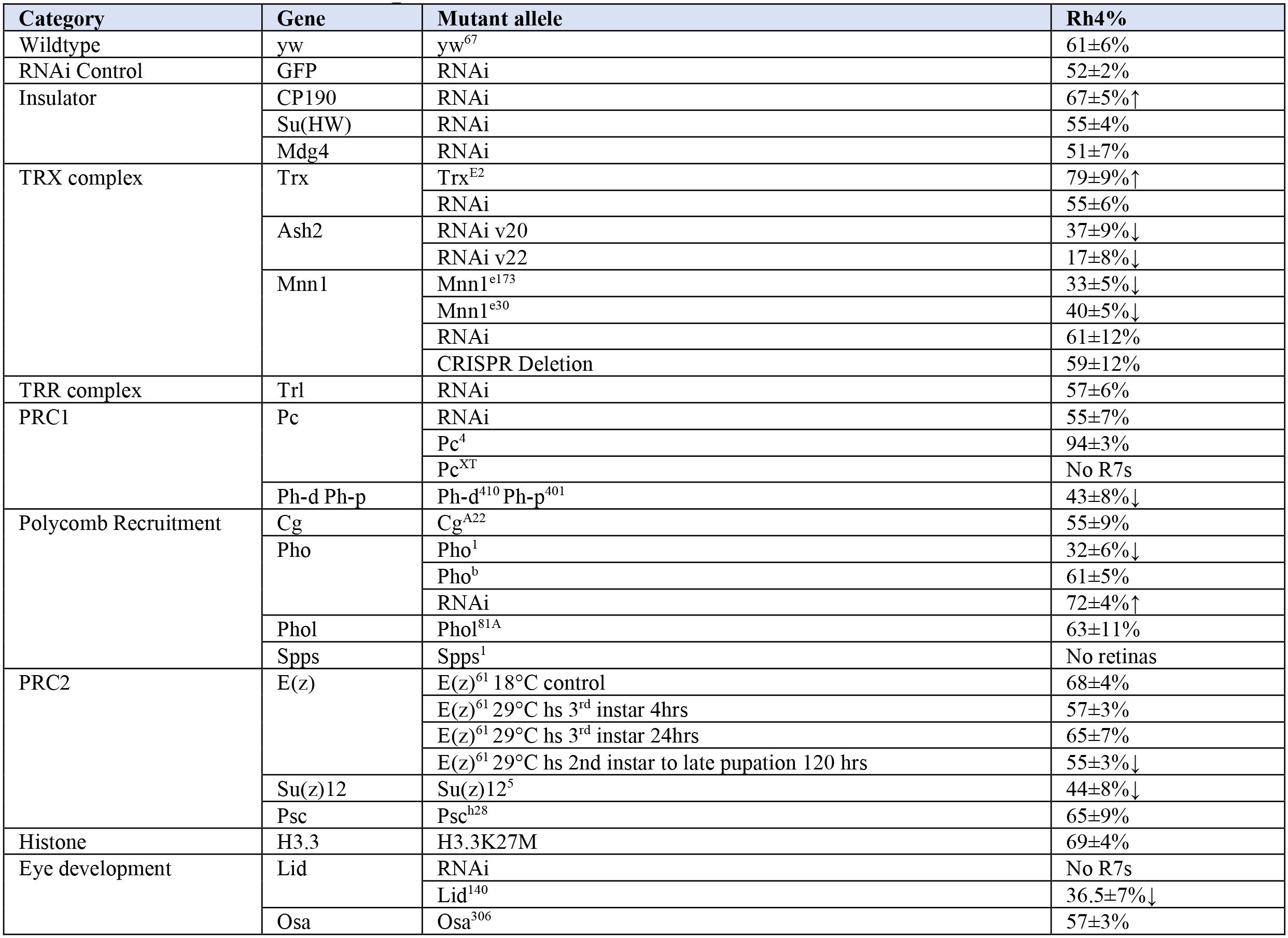
Trans factor regulator screen.

**Table S3.**
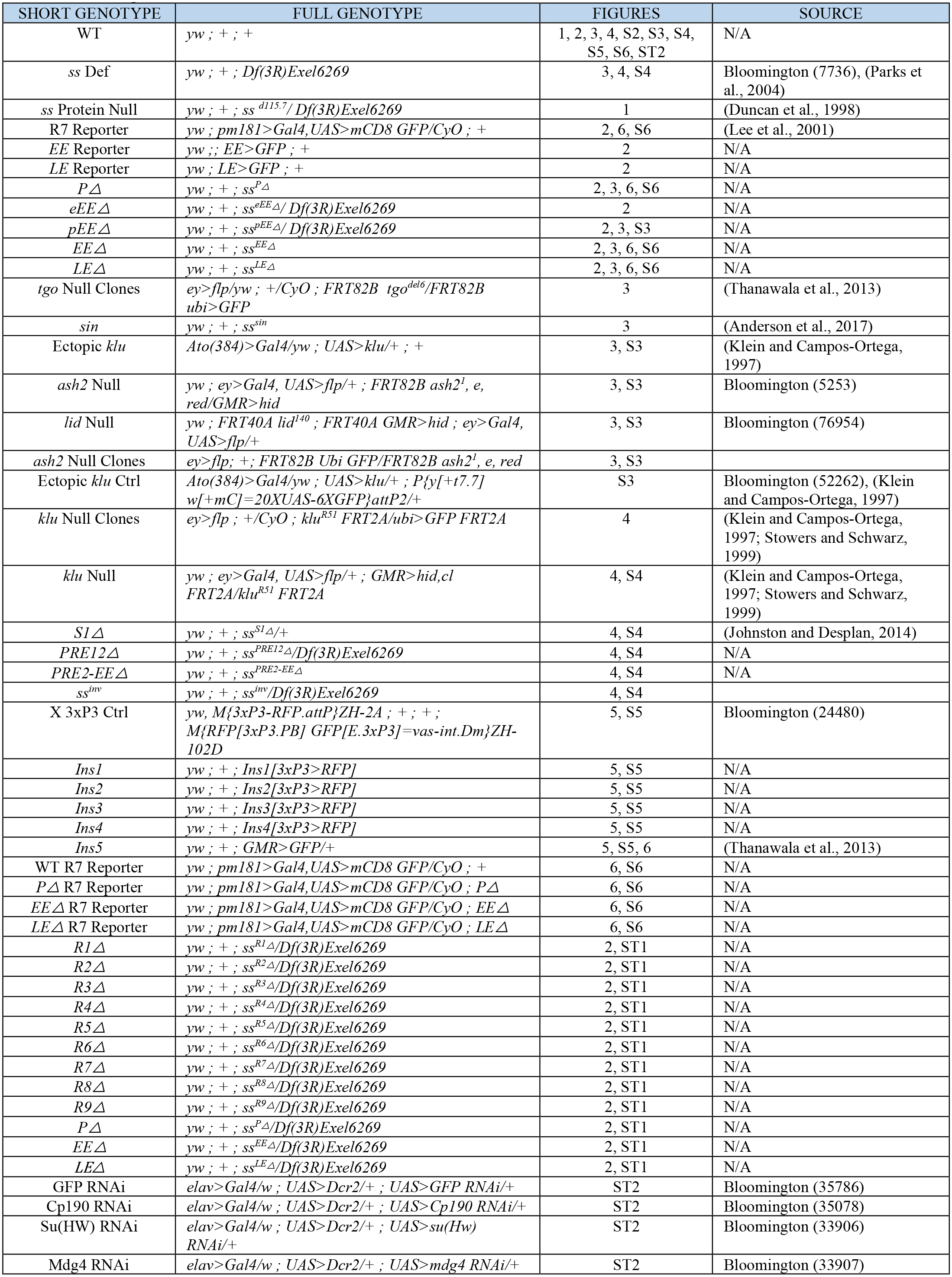
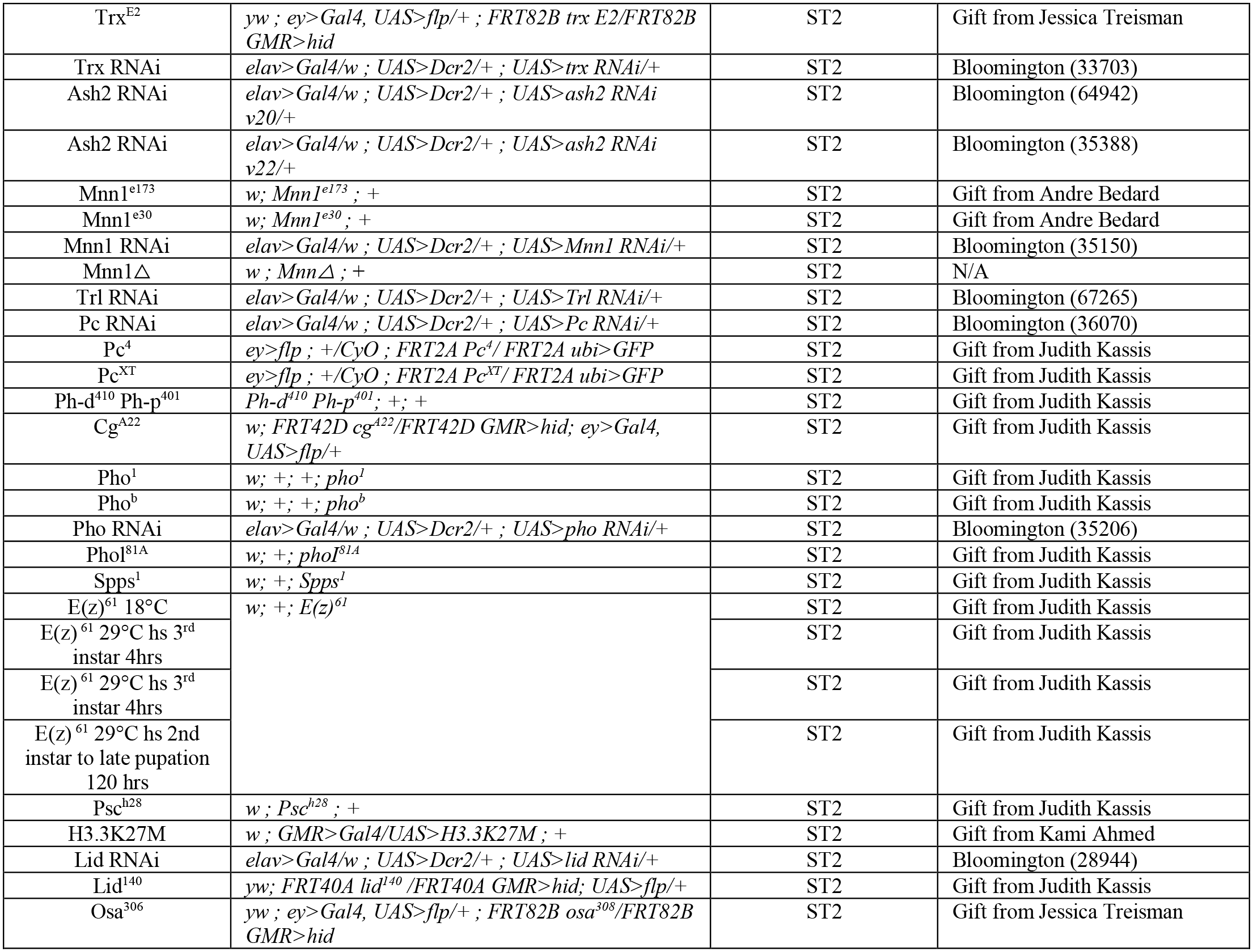
Fly lines.

**Table S4.**
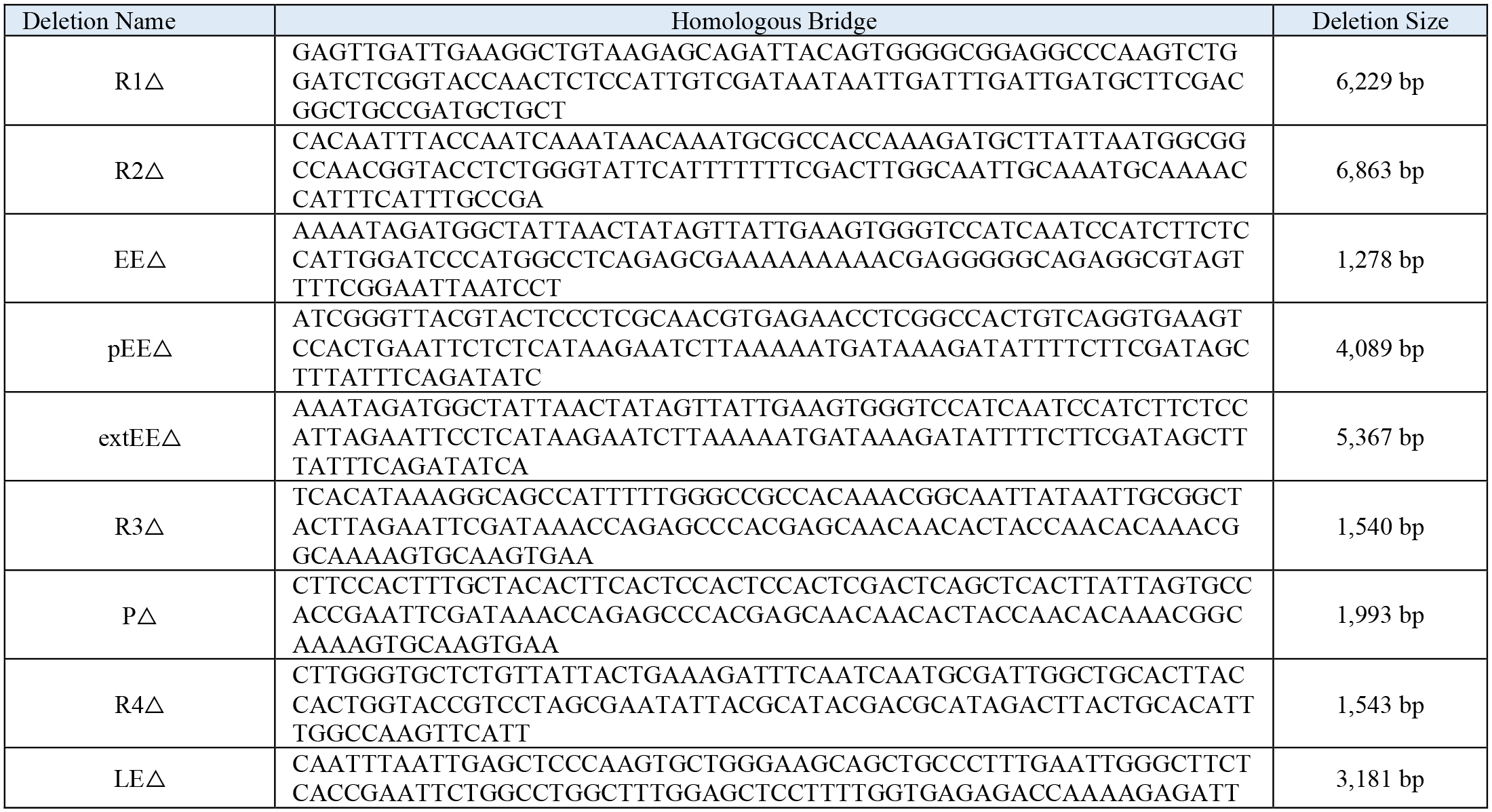
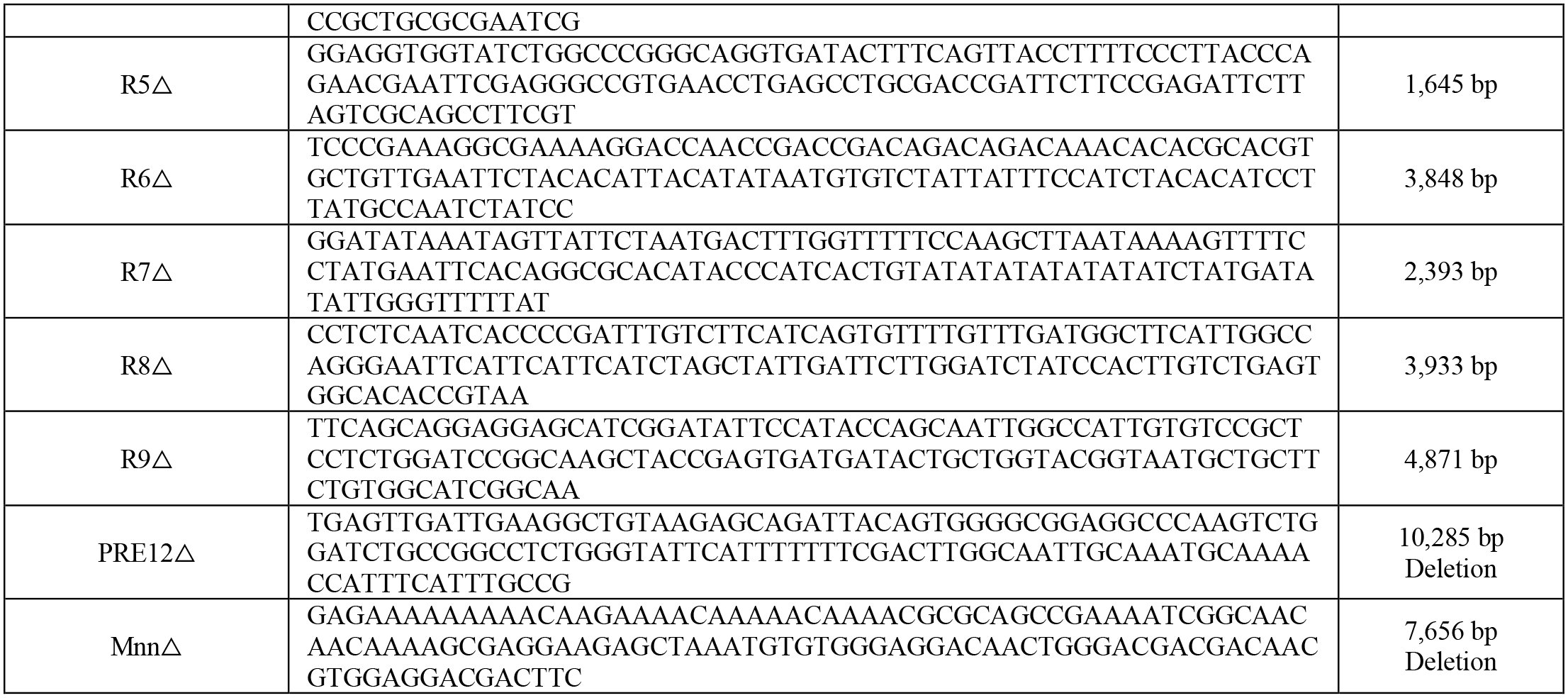
CRISPR Deletion Mutants.

**Table S5.**
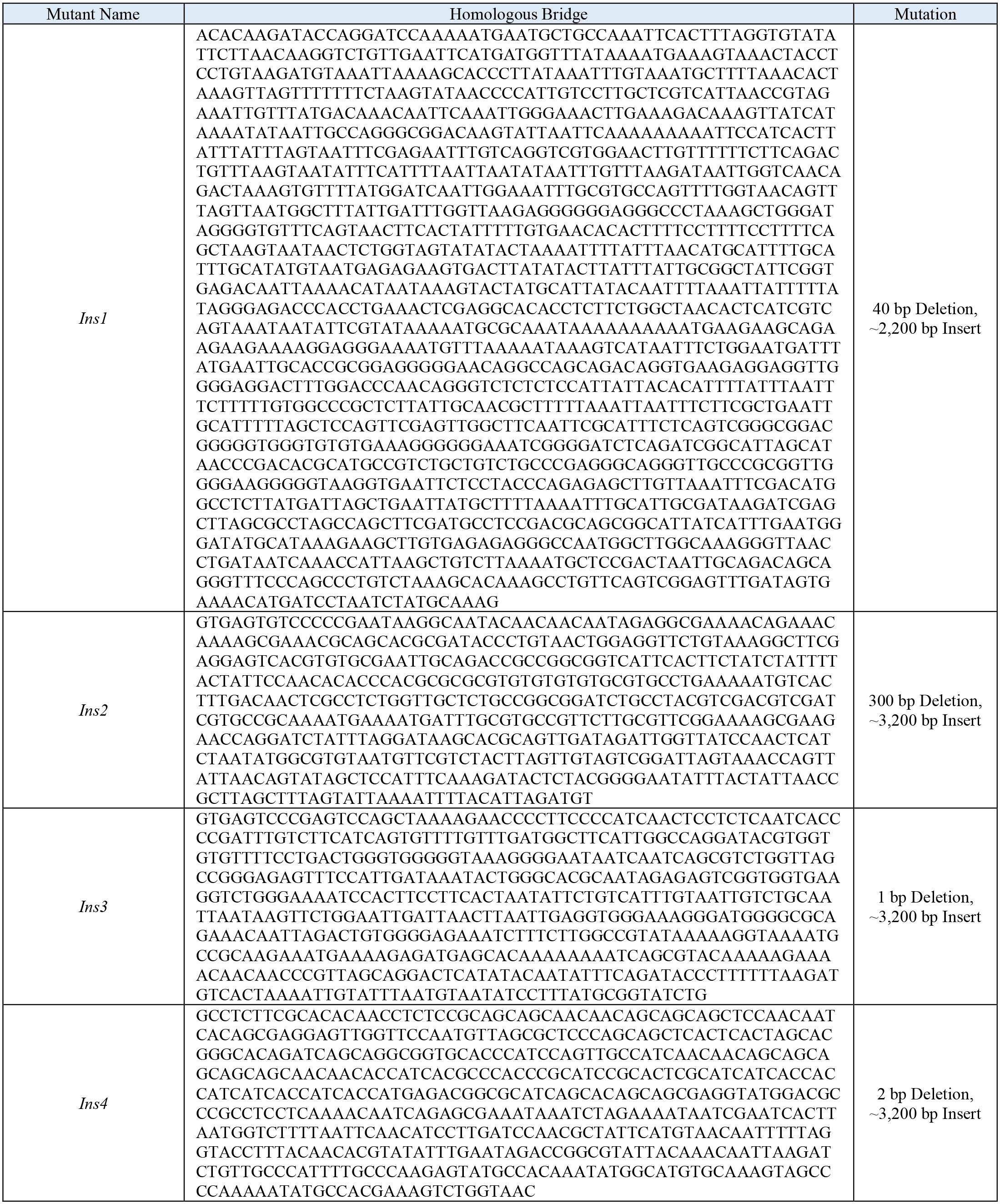
CRISPR Insertion Mutants.

**Table S6.** Oligopaints Probe Libraries.

| Probe Set | Oligo Paints Library | Coordinates | Conjugated Fluorophore |
| --- | --- | --- | --- |
| *ss* RNA | ss-Full-RNA | 3R: 16,370,515-16,435,663 | Cy3 |
| *ss* DNA | ss-Ext-Univ | 3R: 16,370,516-16,435,663 | Cy3 |
| *ss* Upstream DNA | 50-kb extension (left) | 3R: 16,320,533-16,370,533 | Alexa 488 |
| *ss* Downstream DNA | 50-kb extension (right) | 3R: 16,435,681-16,485,681 | Cy5 |

